# Phosphorylation of 53BP1 by ATM enforce neurodevelopmental programs in cortical organoids

**DOI:** 10.1101/2023.05.04.539457

**Authors:** Bitna Lim, Mohamed Nadhir Djekidel, Yurika Matsui, Seunghyun Jung, Zuo-Fei Yuan, Xusheng Wang, Xiaoyang Yang, Abbas Shirinifard Pilehroud, Haitao Pan, Fang Wang, Shondra Pruett-Miller, Kanisha Kavdia, Vishwajeeth Pagala, Yiping Fan, Junmin Peng, Beisi Xu, Jamy C. Peng

**Author notes:** Equal contribution (Bitna Lim, Mohamed Nadhir Djekidel). Present address: Future Medicine Research Institute, Seongnam, Republic of Korea (Bitna Lim). Corresponding author: Jamy C. Peng.

## Abstract

53BP1 is a well-established DNA damage repair factor recently shown to regulate gene expression and critically influence tumor suppression and neural development. For gene regulation, how 53BP1 is regulated remains unclear. Here, we showed that 53BP1-serine 25 phosphorylation by ATM is required for neural progenitor cell proliferation and neuronal differentiation in cortical organoids. 53BP1-serine 25 phosphorylation dynamics controls 53BP1 target genes for neuronal differentiation and function, cellular response to stress, and apoptosis. Beyond 53BP1, ATM is required for phosphorylation of factors in neuronal differentiation, cytoskeleton, p53 regulation, and ATM, BNDF, and WNT signaling pathways for cortical organoid differentiation. Overall, our data suggest that 53BP1 and ATM control key genetic programs required for human cortical development.

## INTRODUCTION

Transcription ensures the appropriate expression of genetic information for the development and function of the organism, whereas DNA repair maintains the integrity of the genetic code. These two processes share cross-functional factors, including CSB, TFII, and XPG, which repair DNA damage generated by torsional stress from transcription-initiating RNA polymerase II (*1–3*). Conversely, some proteins thought to function exclusively in DNA repair have been shown to regulate gene expression. 53BP1 is a key regulator of DNA repair mechanisms, promoting nonhomologous end-joining over homologous recombination (*4*). Upon DNA damage response, 53BP1 (p53 binding protein 1) is required for p53-mediated activation of tumor suppressive genetic programs (*5*). More recently, 53BP1 was found to cooperate with chromatin modifier UTX in neural progenitor cells (NPCs) to promote an open chromatin that facilitates the activation of neurogenic or corticogenic programs (*6*). These findings suggest that gene regulation by 53BP1 is key to tumor suppression and neural development. However, the mechanisms of action of 53BP1 in gene regulation and its upstream mechanism remain to be clarified.

Mechanistic studies of 53BP1 have focused exclusively on the DNA damage response. In order to localize to chromatin at double-stranded breaks, 53BP1 uses its BRCT domain to bind to γH2AX, its Tudor domain to bind to H4K27 dimethylation, and its UDR segment to bind to ubiquitinated H2AK15 (*7–10*). Moreover, the phosphorylated SQ/TQ motif of 53BP1 coordinates the docking of RIF or SCAI to selectively promote nonhomologous end-joining or reduce homologous recombination (*11, 12*). These interactions might be relevant to the gene regulatory activities of 53BP1. For example, γH2AX deposition at transcription start sites is required for R-loop resolution, DNA demethylation, transcription activation, and transcription elongation (*13, 14*).

These studies above have contributed to a general model wherein post-translational modifications of different residues and domains of 53BP1 coordinate its different activities. Most prominently, numerous residues of 53BP1 are phosphorylation by ATM (ataxia telangiectasia mutated) kinase (*10, 15–17*). ATM-mediated phosphorylation of 53BP1 or 53BP1-interacting proteins controls protein interactions, cellular localization, and DNA repair mechanisms (*11, 12, 18–20*). However, whether or how phosphorylation alters the gene regulatory activity of 53BP1 is not known. Here, we uncovered that phosphorylation of 53BP1-serine 25 by ATM and dephosphorylation are required for the appropriate expression of genetic programs during the growth and development of cerebral cortical organoids. ATM-dependent phosphorylation alters neuronal differentiation, cytoskeleton, p53, and ATM, BNDF, and WNT signaling pathways to transcriptionally and post-transcriptionally regulate the development of cortical organoids.

## RESULTS

### Phosphorylated 53BP1-S25 increases during differentiation of hESCs into cortical organoids

Although 53BP1 is necessary for human embryonic stem cells (hESCs) to differentiate into NPCs (*6*), its levels did not change during differentiation (Fig 1A). 53BP1 is targeted by various kinases, including ATM, and we hypothesized that 53BP1 phosphorylation regulates the differentiation of hESCs into NPCs. Intriguingly, we found that the levels of 53BP1 phosphorylated at serine 25 (53BP1-pS25) were markedly increased in NPCs compared to hESCs (Fig 1A, Supplementary Fig 1A–D). The levels of the DNA damage marker γH2AX were similar in NPCs and hESCs (Supplementary Fig 1E), suggesting that the increase in 53BP1-pS25 levels during NPC differentiation is not due to increased DNA damage.

**Figure 1.**
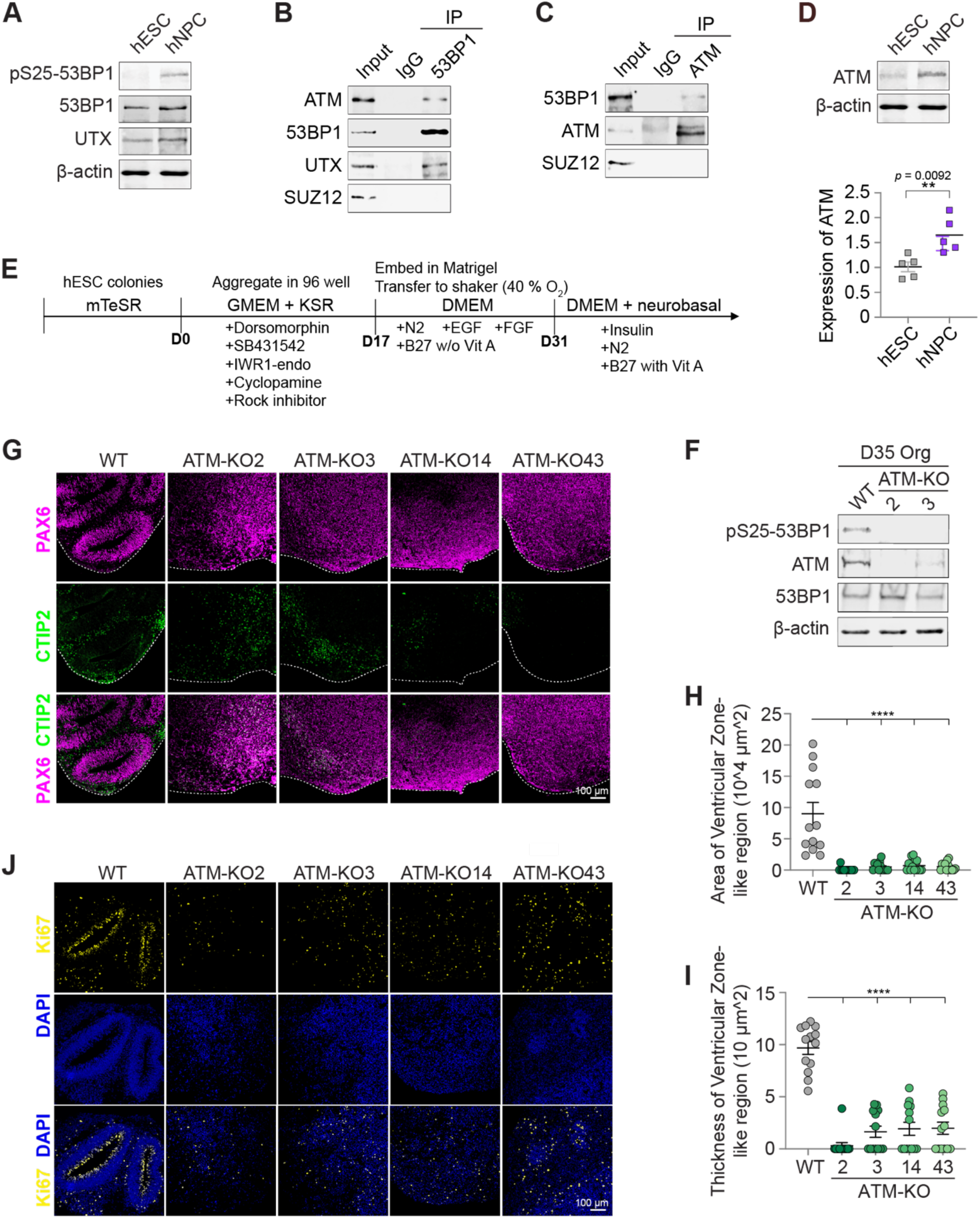
ATM binds 53BP1, is required for pS25-53BP1, and promotes cortical organoid differentiation. WB of the nuclear extract of hESCs and NPCs showed marked increase of 53BP1-pS25 in NPCs. WB analysis of IgG, (B) 53BP1, and (C) ATM co-immunoprecipitation in the nuclear extract of hESCs. (D) Quantification of the relative ATM protein levels in 5 replicate WB analyses of hESCs and NPCs. (E) Schematic diagram of the cortical organoid differentiation. Aggregates were formed in the induction media for 17 days, embedded in Matrigel droplets and cultured in cortical differentiation medium for 16 days, and then cultured in cortical maturation media thereafter. GMEM, Glasgow Modified Essential Medium. KSR, Knockout Serum Replacement. DMEM, Dulbecco’s Modified Eagle Medium. (F) WB analysis of WT and ATM-KO cortical organoids at day 35 of differentiation. Immunofluorescence of (G) PAX6 and CTIP2 and (J) KI67 in cryosections of cortical organoids at day 35 of differentiation. Bar, 100 μm. At day 35 of differentiation, the (H) area and (I) thickness of ventricular zone (VZ)-like regions were compared between groups. Data points represent single organoids. The mean ± SEM values were compared by one-way ANOVA with Dunnett’s multiple comparisons test to yield **** indicating p< 0.0001. n= 13 organoids/group.

### ATM is required for 53BP1-S25 phosphorylation and hESC differentiation into cortical organoids

The ATM kinase phosphorylates 53BP1-S25 (*15*), and we thus investigated whether ATM plays a role in neural differentiation. First, we found that ATM co-immunoprecipitated with 53BP1, as did the positive control UTX, but not with the negative control SUZ12 (core subunit of PRC2, which does not bind these proteins; Fig 1B). Similarly, 53BP1 co-immunoprecipitated with ATM, but not with the negative control SUZ12 (Fig 1C). Like 53BP1-pS25, ATM levels were significantly increased in NPCs compared with hESCs (Fig 1D). ATM upregulation in NPCs was shown by a previous DNA damage response study (*21*).

Next, we used the CRISPR-Cas9 system to generate four *ATM*-knockout (KO) hESC lines (Supplementary Fig 1F, 2A–B). Data from RNA-seq and immunofluorescence showed that *ATM*-KO did not markedly alter hESC pluripotency (Supplementary Fig 2C–D). To analyze the role of ATM in human cortical development, we used an established protocol to differentiate WT and *ATM*-KO hESCs into cortical organoids (Fig 1E, Methods, (*6*)). We did not detect 53BP1-pS25 in *ATM*-KO D35 cortical organoids (Fig 1F) nor NPCs (Supplementary Fig 2B), consistent with loss of ATM-mediated phosphorylation of 53BP1-S25 during neural differentiation of hESCs. *ATM*-KO modestly reduced γH2AX levels in NPCs (Supplementary Fig 2E), suggesting that ATM promotes the phosphorylation of H2AX-S139 in NPCs.

By day 35 (D35) of differentiation, WT cortical organoids expressed the forebrain NPC marker PAX6 in ventricular zone-like regions that were surrounded by cells expressing the subcortical projection neuron marker CTIP2 (Fig 1G). In contrast, *ATM*-KO D35 cortical organoids displayed disorganized ventricular zone-like regions, fewer cells positive for the proliferation marker KI67, and reduced neuronal differentiation (Fig 1G–J). By D55, a subset of *ATM*-KO cortical organoids appeared smaller than controls, but this difference was not statistically significant (Supplementary Fig 3A–B). Thus, ATM is required for both NPC proliferation and neuronal differentiation in cortical organoids.

### ATM safeguards transcriptional and translational programs in differentiating cortical organoids

We used RNA-seq to profile the transcriptomes of D35 cortical organoids derived from WT and *ATM*-KO hESCs. Using a false discovery rate (FDR) <0.05, we identified 590 upregulated genes and 1,084 downregulated genes in 8 *ATM*-KO versus 6 WT datasets. Upregulated genes were highly enriched in terms related to cellular metabolism pathways (e.g. ribosome biogenesis, rRNA processing, spliceosome complex, post-transcriptional gene regulation, regulation of translation; Fig 2A), whereas downregulated genes were highly enriched in terms related to neuronal differentiation (e.g. neuron projection guidance, postsynaptic membrane; Fig 2B). These data suggest that ATM restricts metabolism and promotes neuronal differentiation in the cortical organoids.

**Figure 2.**
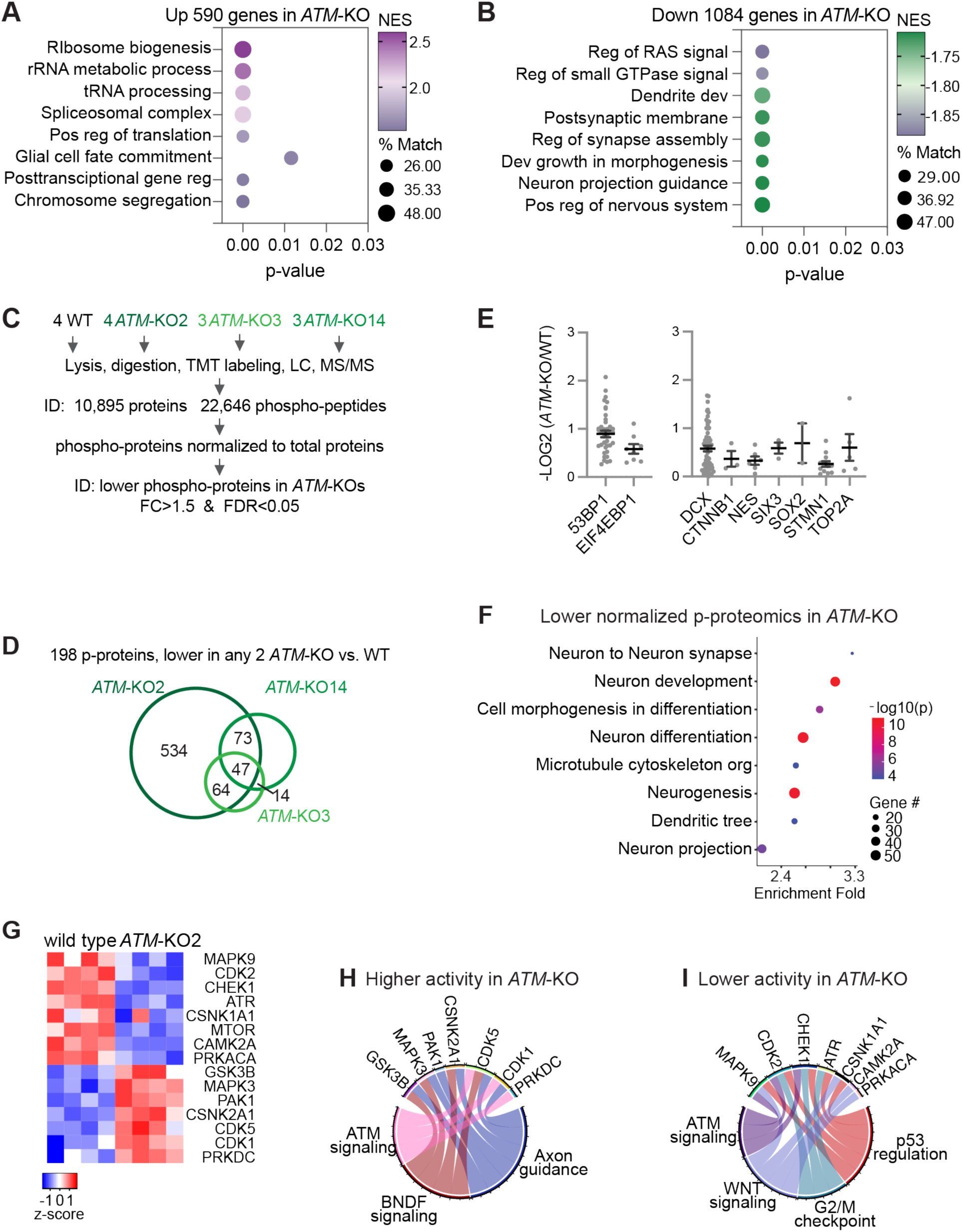
Transcriptomic and proteomic profiles of WT versus *ATM*-KO cortical organoids. From, gene set enrichment analysis, functional terms that are highly enriched in (A) upregulated and (B) downregulated genes in *ATM*-KO D35 cortical organoids. NES, normalized enrichment score. % Match, % of genes in the enriched term that overlap the differentially expressed genes or proteins. (C) Schematic diagram outlining TMT LC-MS/MS profiling of total proteomics and phospho-proteomics of D35 WT and *ATM*-KO cortical organoids. TMT signals from total proteomics were used to normalize those of phospho-peptides. (D) Using, FC>1.5 and FDR<0.05, 198 phospho-proteins were found to be lower in 2 *ATM*-KO versus WT. (E) Normalized levels of phospho-proteins that have ATM-dependent phosphorylation in D35 cortical organoids. 53BP1 and EIF4EBP1 were known substrates of ATM. (F) Enrichment of proteins with ATM-dependent phosphorylation in specific functional categories. (G) Heatmap showing altered activities of kinases between D35 *ATM*-KO2 and WT cortical organoids. Relative changes in kinase activity are shown as row Z-scores. Kinase activity was inferred by IKAP (*24*) based on normalized substrate phosphorylation levels from phosphor-proteome. The normalization was performed by dividing phosphor-peptide abundance of each protein by corresponding protein abundance (*52*). Circos plots showing kinases with inferred (H) higher and (I) lower activities in D35 *ATM*-KO versus WT cortical organoids and their corresponding enriched pathways.

We used multiplexed tandem mass tag-based quantification and 2-dimensional liquid chromatography-tandem mass spectrometry (TMT LC-MS/MS) to profile the proteome and phosphoproteome of WT and *ATM*-KO D35 cortical organoids (Supplementary Fig 3C, Methods). We quantified 10,895 proteins between 4 WT, 4 *ATM*-KO2, 3 *ATM*-KO3, and 3 *ATM*-KO14 D35 cortical organoid samples by using the criteria of fold change >1.5 and FDR<0.05 (Fig 2C). Consistency between replicate datasets is supported by principal component analysis (Supplementary Fig 3C). Gene set enrichment analysis (GSEA) showed that compared to WT, upregulated proteins in *ATM*-KO were enriched in terms related to neurotransmission, neuron spine, dendrite, synapse, and axon (Supplementary Fig 3D), whereas downregulated proteins were enriched in BMP/TGFβ and WNT signaling, epithelial morphogenesis, and stem cell differentiation (Supplementary Fig 3E). These data suggest that ATM controls post-transcriptional and translational gene regulation to suppress neuronal function and promote stem cell differentiation, epithelial morphogenesis, and TGFβ and WNT signaling pathways in D35 cortical organoids.

We note that the transcriptomics and proteomics data show different patterns in the ATM-KO. While transcriptomic programs related to neuronal differentiation were downregulated (Fig 2B), proteomic programs related to neuronal function were upregulated (supplementary Fig 3D) in *ATM*-KO versus WT cortical organoids. We surmised that ATM regulates stages of neuronal specification, differentiation, and maturation. Dysregulated transcriptomic and proteomic programs were consequences of the pleiotropic effect of ATM.

### ATM-dependent phosphorylation controls signaling pathways for neurogenesis, stem cell differentiation, and morphogenesis in cortical organoids

To investigate how ATM exerts its modulatory control during cortical organoid formation, we performed phospho-proteomics analysis of WT and ATM-KO cortical organoids. Using TMT LC-MS/MS, we quantified 22,646 phospho-peptides and normalized their abundance based on the protein abundance measured in the total proteomics analysis. Compared with WT, 198 phosphorylated proteins were consistently lower in at least 2 of the 3 *ATM*-KO lines (log2(fold change >1.5) and FDR<0.05; Fig 2D, Supplementary Table 1). Amongst these proteins, 53BP1 and EIF4EBP1 were known substrates of ATM (Fig 2E) (*15, 16, 22, 23*), validating our approach to identify putative ATM substrates in cortical organoids. We note that this approach does not distinguish between direct and indirect effects, and thus some of those proteins could be phosphorylated by protein kinases that require ATM for their activity. Intriguingly, many ATM-dependent phosphorylated proteins were key neurodevelopmental regulators (Fig 2E) and enriched in functions in neurodevelopment, neurogenesis, cell morphogenesis, and cytoskeleton (Fig 2F).

We also found 172 proteins that had consistently higher levels of phosphorylation in any 2 *ATM*-KO compared to WT. These proteins were enriched in nonhomologous end joining, telomere maintenance, cytoskeleton, and synapse (Supplementary Fig 3E). These data suggest that ATM restricts downstream kinase activity over substrates involved in these pathways. To explore the downstream protein kinases that could be controlled by ATM, we used the IKAP machine learning algorithm (*24*) to analyze substrates (inferred from literature curation) and deduce the activities of those kinases. For example, in *ATM*-KO compared to WT, we found reduced phosphorylation of DCX, MAPT, and NFATC4 (Supplementary Fig 4A, Table 2), which were related to MAPK9 activities by IKAP (*24*). In contrast, we found higher phosphorylation of ADD2, ADD3, DCX, DNM1L, DPYSL3, MAPT, and SRC (Supplementary Fig 4B, Table 2), which are known were related to CDK5 activities by IKAP (*24*). In *ATM*-KO, we inferred lower activities in MAPK9, CDK2, CHEK1, ATR, CSNK1A1, MTOR, CAMK2A, and PRKACA (FIG 2G, Supplementary Fig 4C), with enriched function in ATM signaling, BNDF signaling, and axon guidance (FIG 2G–H, Supplementary Fig 4C). On the other hand, we inferred higher activities in GSK3B, MAPK3, PAK1, CSNK2A1, CDK5, CDK1, and PRKDC (FIG 2G, I, Supplementary Fig 4C), with enriched function in ATM signaling, WNT signaling, G2/M checkpoint, and p53 regulation in *ATM*-KO (Fig 2H–I). Of these kinases, CHEK1, ATR, and PRKDC were known substrates of ATM (*23, 25*); thus, some of the altered kinase activities could be secondary to *ATM*-KO. Collectively, these data support that kinases for ATM signaling, BNDF signaling, WNT signaling, G2/M checkpoint, and p53 became dysregulated in *ATM*-KO D35 cortical organoids.

We thus conclude that ATM controls key neurodevelopmental regulators and exerts strong and pleiotropic influences over the differentiation of cortical organoids. In *ATM*-KO, dysregulated phosphorylation and activities of those key neurodevelopmental regulators disrupt the transcriptomic program for neuronal differentiation and result in higher proteomic programs for neuronal function. These dysregulated programs would lead to the defects in neurogenesis and morphogenesis observed in D35 *ATM*-KO cortical organoids.

### Phosphorylation of 53BP1-S25 is required for NPC proliferation

We next examined ATM-dependent phosphorylation of 53BP1-S25. To specifically investigate the functional significance of 53BP1-pS25, we used the CRISPR-Cas9 system to mutate the endogenous 53BP1 serine 25 to alanine (S25A) or aspartic acid (S25D) (Supplementary Fig 4D, Fig 3A, Methods). The alanine substitution precludes phosphorylation, whereas aspartic acid is chemically similar to phosphoserine (*26*). We generated four 53BP1-S25A hESC lines (34-3, 34-4, 79-1, and 79-3) and four 53BP1-S25D hESC lines (14-3, 14-15, 14-19, and 17). The total levels of 53BP1 were similar in WT, 53BP1-S25A, and 53BP1-S25D NPCs, and we did not detect pS25 in 53BP1-S25A NPCs, as expected (Supplementary Fig 1D and 4E). Control (mock transfected WT), 53BP1-S25A, and 53BP1-S25D hESC lines displayed similar transcriptomic profiles and pluripotency marker expression (Supplementary Fig 2C, 4F, 5A), suggesting that 53BP1-S25A and –S25D do not affect hESC self-renewal.

**Figure 3.**
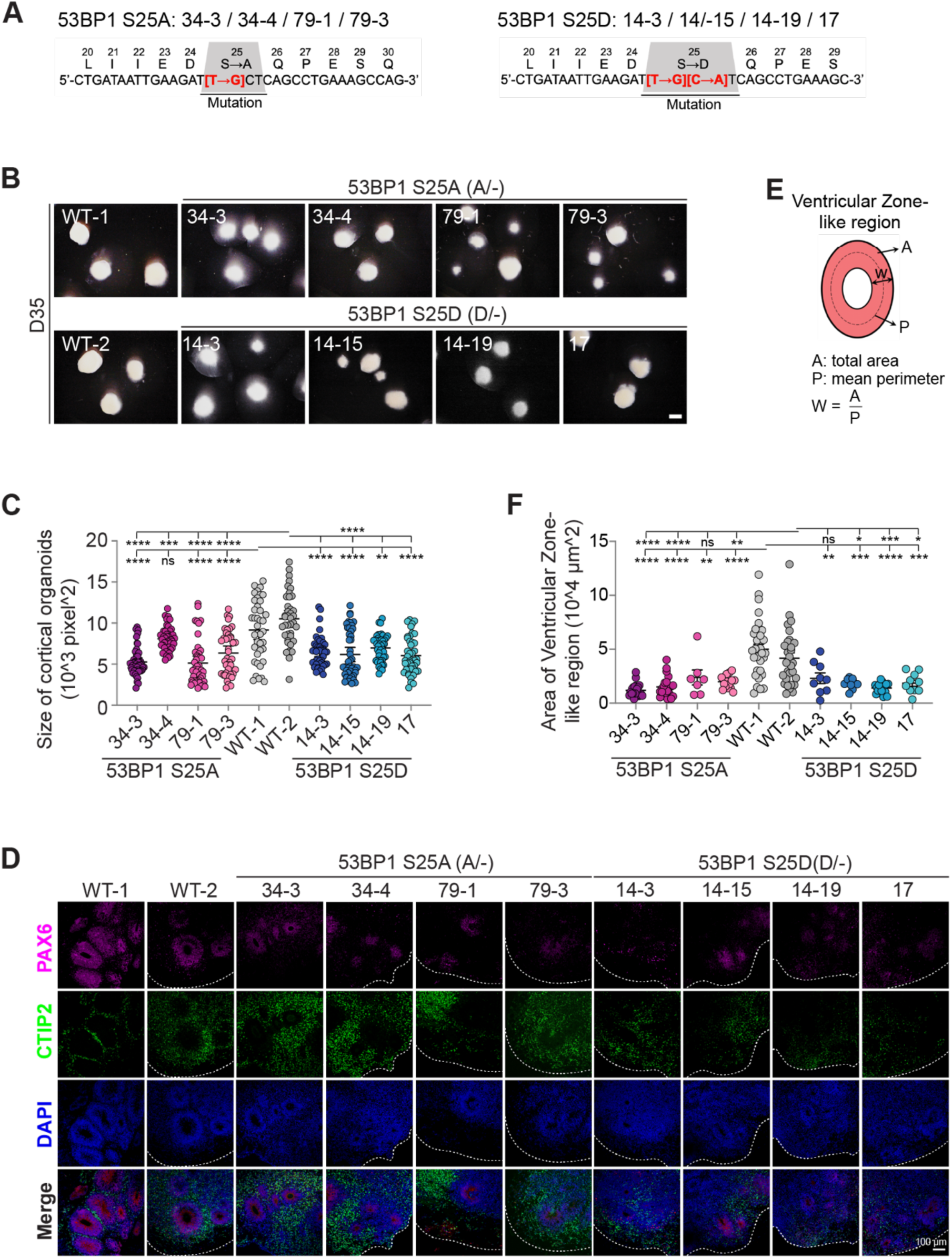
53BP1-pS25 is required for the differentiation of cortical organoids. (A) In the endogenous *53BP1* locus, the codon TCT encoding serine-25 in was mutated to GCT and GAT encoding alanine and glutamate, respectively. (B) Bright-field images of cortical organoids formed by 4 53BP1-S25A lines, 4 53BP1-S25D lines, and 2 WT control at day 35 of differentiation. Bar, 1.5 mm. At day 35 of differentiation, the (C) organoid size and (F) area of ventricular zone-like region were compared between groups. Data points represent single organoids. The mean ± SEM values were compared by one-way ANOVA with Dunnett’s multiple comparisons test to yield ****, ***, **, *, and ns indicating p< 0.0001, 0.001, 0.01, 0.05, and not significant, respectively. n = 39 – 47 organoids/group for (C) and 15 – 33 organoids/group for (F). (D) Immunofluorescence of PAX6 and CTIP2 in cryosections of cortical organoids at day 35 of differentiation. Bar, 100 μm. (E) Illustration of ventricular zone-like areas in cortical organoids.

**Figure 4.**
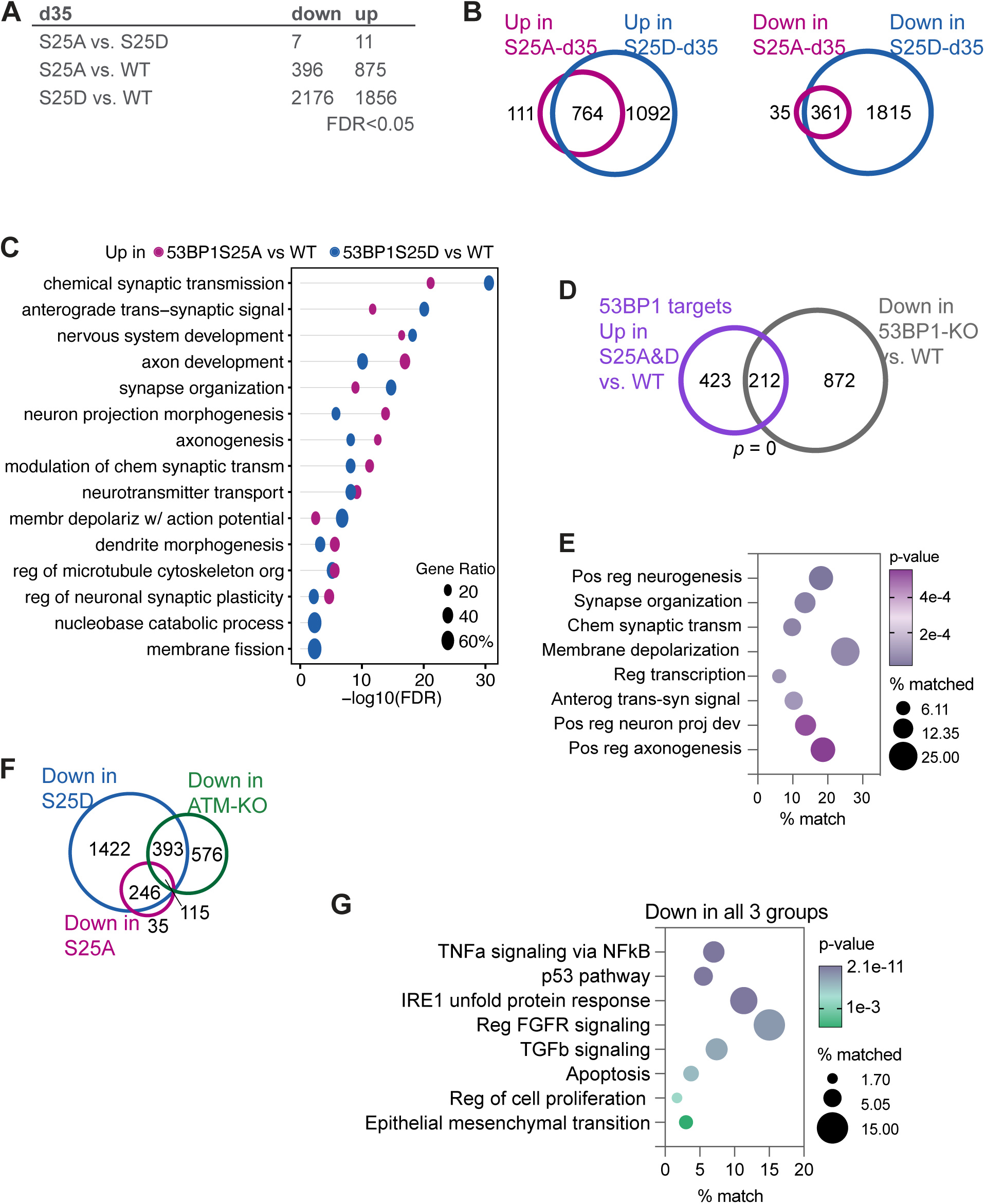
53BP1-S25 phosphorylation enforce the appropriate expression of genetic programs for cortical organoid differentiation. (A) Number of differentially expressed genes identified by pair-wise comparisons at FDR <0.05. At day 35 of differentiation, 53BP1-S25A and S25D cortical organoids are molecularly similar. (B) Differentially expressed genes in 53BP1-S25D versus WT overlap 87% (764/875) and 91% (361/396) of those in 53BP1-S25A versus WT. (C) Extensive overlap of upregulated GSEA terms between 53BP1-S25A versus WT and 53BP1-S25D versus WT. Most terms relate to axon, synapse, and neurotransmitter. (D) Of 53BP1 target genes upregulated by S25A and S25D, 212 genes require WT 53BP1 for expression in cortical organoids. (E) The 212 genes are enriched in functions related to transcriptional regulation, neuron projection, axonogenesis, synapse, neurotransmitter synthesis and transport, and membrane depolarization. (F) Venn diagrams depict high overlaps between downregulated genes in all 3 groups of mutant versus WT pairwise comparisons. (G) GSEA terms of the 115 genes that were downregulated in all 3 groups (versus WT) revealed the genetic programs co-promoted by ATM and 53BP1-pS25.

To analyze the role of 53BP1-S25 in human cortical development, we differentiated control WT, 53BP1-S25A, and 53BP1-S25D hESCs into cortical organoids (Fig 1E, Methods). The 53BP1-S25A and –S25D D35 cortical organoids were smaller than controls (Fig 3B–C) and had significantly smaller ventricular zone-like regions (Fig 3D–F), suggesting they had fewer NPCs. Additionally, we observed fewer cells that were positive for Ki67 (proliferation marker) or for phosphorylated-serine 10 histone H3 (mitotic chromatin marker) in 53BP1-S25A and –S25D D35 cortical organoids compared to WT (Supplementary Fig 5B–D). The proportion of CTIP2-positive subcortical projection neurons were similar between WT, 53BP1-S25A and –S25D D35 cortical organoids (Supplementary Fig 6A). At D55, the 53BP1-S25A and S25D cortical organoids remained significantly smaller than WT (Supplementary Fig 6B–D, Table 3-4). Overall, these data suggest that 53BP1-S25A and S25D had similar cell biological effects and that phosphorylation of 53BP1-S25 is required for proliferation of NPCs and growth of cortical organoids.

### Phosphorylation of 53BP1-S25 modulates the expression of genetic programs for neuronal differentiation and function

Using RNA-seq, we examined the transcriptomes of WT (6 samples), 53BP1-S25A (8 samples), and 53BP1-S25D D35 (8 samples) D35 cortical organoids. We analyzed expressed genes with count per million values >1 and observed few differences in gene expression between 53BP1-S25A and 53BP1-S25D D35 cortical organoids, using FDR <0.05 (Fig 4A). Separate comparisons of 53BP1-S25A versus WT and 53BP1-S25D versus WT revealed high concordant changes in gene expression caused by the 2 mutations, as >87% of differentially expressed genes in 53BP1-S25A were also altered in 53BP1-S25D (Fig 4B and Supplementary Fig 7A). However, 53BP1-S25D disrupted the expression of 2-3 fold more genes than 53BP1-S25A, suggesting that the 53BP1-S25D mutation led to gain-of-function. To explore this further, we performed GSEA and found high overlap of the top terms in the upregulated genes in 53BP1-S25A versus WT and 53BP1-S25D versus WT (Fig 4C). In contrast, there was low overlap of the top terms in the downregulated genes in 53BP1-S25A versus WT and 53BP1-S25D versus WT (Supplementary Fig 7B). Intriguingly, genes upregulated by 53BP1-S25A and S25D specifically relate to synapse, axon, and neurotransmitter (Fig 4C). These were the same categories enriched in genes upregulated by 53BP1-S25D but not S25A (Supplementary Fig 7D). Thus, compared to 53BP1-S25A, S25D upregulates more genes involved in neuronal function.

To dig deeper into this intriguing observation, we compared our data with previous published transcriptomic data that compared *53BP1*-KO and WT cortical organoids, which support a requirement of 53BP1 for activating neurogenic genes (*6*). We had noticed that GSEA categories of genes upregulated by 53BP1-S25A and S25D were similar to those downregulated by *53BP1*-KO. This comparison showed a significant overlap of 212 genes upregulated by 53BP1-S25A and –S25D with 53BP1-bound target genes that were downregulated in *53BP1*-KO versus WT (*p* = 0 by empirical estimation, Fig 4D). As the proportion of CTIP2-positive subcortical projection neurons did not differ between WT, 53BP1-S25A, and 53BP1-S25D (Supplementary Fig 6A), the transcriptomic changes likely reflect higher expression of neuronal transcripts in 53BP1-S25A and S25D neurons. The 212 genes were enriched in regulation of transcription, neurogenesis, neuronal projection, axonogenesis, synapses organization, and membrane depolarization (Fig 4E). This overlap suggests that in cortical organoids, these 212 genes require and are upregulated by 53BP1-pS25.

How do transcriptomic changes in 53BP1-S25A and S25D compared to those in *ATM*-KO cortical organoid? We observed little overlap between the downregulated genes in *ATM*-KO and the upregulated genes in 53BP1-S25A and –S25D. In contrast, we observed greater overlap in concordant gene expression changes in *ATM*-KO, 53BP1-S25A, and 53BP1-S25D versus WT (Fig 4F, Supplementary Fig 6C–D). Notably, all 3 mutant types shared downregulated genes that were enriched in TNFα signaling via NFKB, p53 pathway, IRE1-mediated unfolded protein response, FGFR signaling, TGFý signaling, apoptosis, regulation of cell proliferation, and epithelial mesenchymal transition (Fig 4G). These data suggest that both ATM and 53BP1-pS25 promote the expression of these genes. We inferred that a role of ATM is to promote the expression of these genes via phosphorylating 53BP1-S25 in D35 cortical organoids.

### 53BP1-S25A and S25D predominantly alter the expression of 53BP1 target genes

To obtain further mechanistic insights into the role of 53BP1 in controlling gene expression, we reanalyzed 53BP1 ChIP-seq data (using 2 separate anti-53BP1 antibodies) in NPCs (*6*) using a more stringent statistical threshold. Using SICER (*27*) and MACS2 (*28*) with FDR <0.05, we identified 37,519 targets bound by 53BP1. About 41% of the 53BP1 targets localize to promoter regions, 76.7% in gene bodies, and 23.3% were intergenic (Supplementary Fig 7E). This enrichment of 53BP1 at promoters suggest a transcriptional regulatory role of 53BP1. More than 82% of the differentially expressed genes in 53BP1-S25A and –S25D D35 cortical organoids were targets bound by 53BP1 (Fig 5A–C). 53BP1 target genes with increased transcript levels in the mutant organoids were highly enriched in neuronal development, axonogenesis, neuron projection, synapse organization, and neurotransmitter transport, transmission, and signaling (Fig 5D), whereas 53BP1 targets with reduced transcript levels in the mutant organoids were enriched in IRE1-mediated unfolded protein response, cellular response to stress, iron import, and apoptosis regulation (Fig 5E). Therefore, 53BP1-pS25 directly controls the expression of neuronal function programs. In particular, we note that genes involved in IRE1-mediated unfolded protein response and apoptosis regulation had reduced expression upon loss of ATM or mutation of 53BP1-S25, and are direct targets of 53BP1 in NPCs. These data suggest that ATM-mediated phosphorylation of 53BP1-S25 directly promotes the expression of these genes to maintain NPCs during formation of cortical organoids.

**Figure 5.**
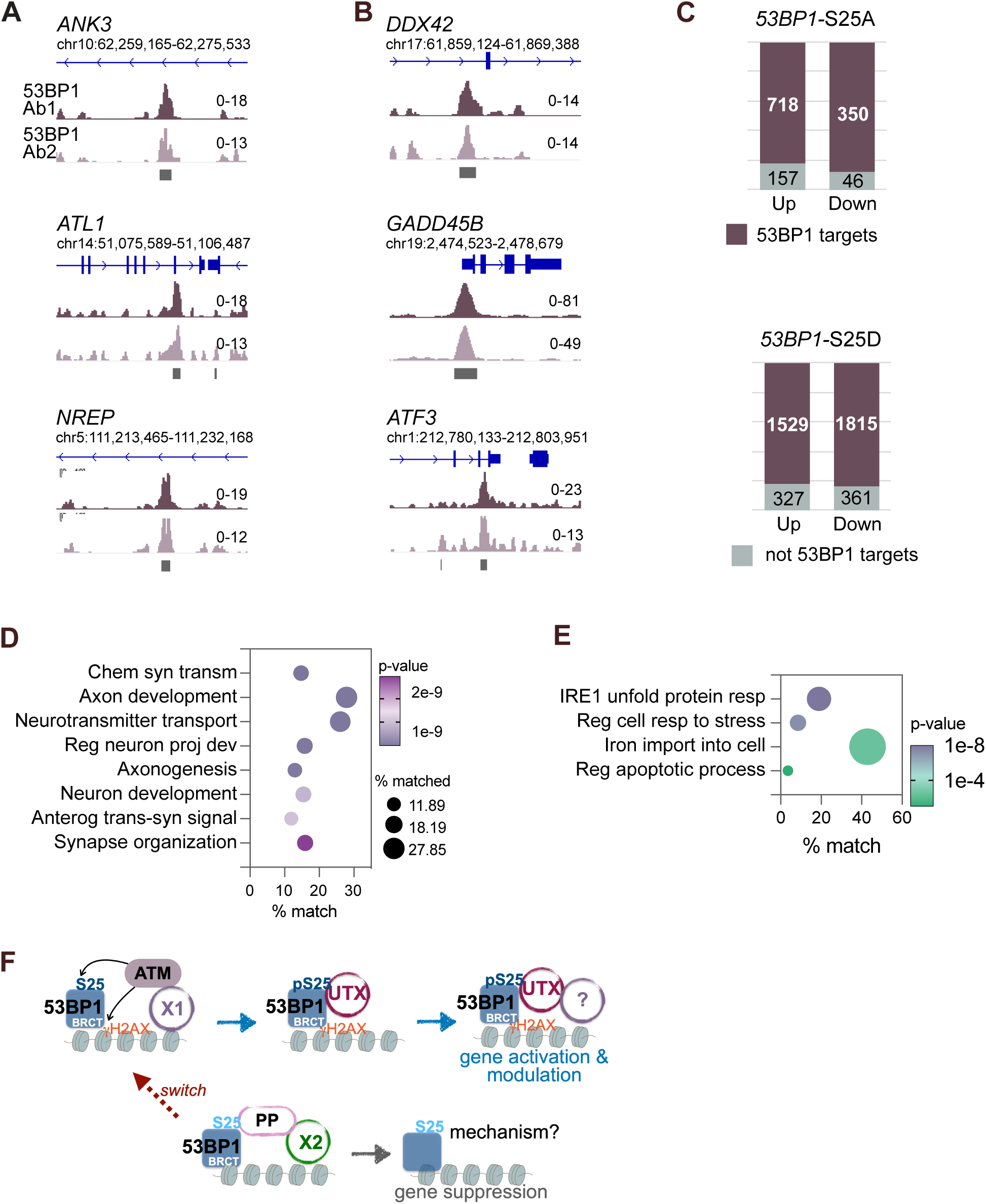
53BP1-pS25 positively and negatively regulate 53BP1 target genes. 53BP1 ChIP-seq tracks at loci of representative (A) upregulated and (B) downregulated genes in 53BP1-S25A and S25D versus WT D35 cortical organoids. (C) More than 82% of differentially expressed genes in 53BP1-S25A or S25D versus WT are targets of 53BP1. (D) 53BP1-S25A and S25D upregulate 53BP1 targets that are involved in neuron development and projection, axonogenesis, synapse, and neurotransmitter synthesis and transport. (E) S25A and S25D downregulate 53BP1 targets that are enriched in IRE1-mediated unfolded protein response, regulation of cellular response to stress, iron import into cells, and regulation of apoptosis. (F) In the proposed model, an unknown factor recruits ATM, which phosphorylates H2AX, which in turn binds to the BRCT domain of 53BP1. 53BP1 is phosphorylated by ATM and interacts with UTX and other factors to promote open chromatin structure and gene activation then modulation. For suppressed genes, an unknown factor leads to the dephosphorylation of γH2AX and 53BP1. Via a switch mechanism, the suppressed genes become activated.

## DISCUSSION

We report that ATM exerts strong pleiotropic influences over transcriptional, post-transcriptional, and translational control of gene expression during the differentiation of hESCs to cortical organoids. Our in vitro model does not fully recapitulate neurodevelopment in vivo, but it may yield information significant to corticogenesis. We showed that ATM-dependent phosphorylation predominantly influences factors in neurogenesis, neuronal differentiation, cell morphogenesis, and microtubule cytoskeleton as well as kinases involved in ATM, BNDF, and WNT signaling, G2/M checkpoint, and p53 regulation. These findings have broad implications about diseases associated with ATM, including ataxia telangiectasia (*29–31*).

Because of the pleiotropic effect of ATM, we focused our studies on 53BP1-pS25, which depends on ATM and regulates genetic programs including signaling pathways, p53 regulation, apoptosis, and cell proliferation. Current understanding about 53BP1 mechanisms in DNA damage response helps us form a model about ATM–53BP1 mechanism in gene regulation (Fig 5F). In our model, ATM phosphorylates H2AX at transcription start sites (*13, 14*) to recruit 53BP1 and phosphorylate 53BP1-S25. The fidelity of gene expression requires 53BP1-pS25 dynamics and likely involves 53BP1 interactors such as RIF, SCAI, and UTX (*6, 11, 12*) or other transcriptional regulators in the 53BP1 network (*32*).

Our findings suggest that regulation of neurodevelopmental programs by ATM–53BP1 is remarkable for (a) maintaining NPCs and the size of cortical organoids, (b) driving and modulating programs for synapse, axon, and neurotransmitter, and (c) involving a temporal component. As cortical organoids progress in differentiation, the temporal coordination of initial suppression of neuronal differentiation and function in NPCs followed by 53BP1-mediated activation and modulation of the same programs requires a switch (Fig 5F). This switch involves ATM and 53BP1-pS25 dynamics and specifically controls genes involved in synapse, axon, and neurotransmitter (Fig 4C), which are crucial to cognition. In the future, elucidation of this mechanism will shed light on the molecular control of corticogenesis. Beyond 53BP1, ATM-dependent phosphorylation likely controls many key neurodevelopmental regulators. Future studies of how ATM selects substrates to exert its pleiotropic influences will advance our understanding of epigenetic programming of human neurodevelopment.

## ACKNOWLEDGEMENTS

The authors thank A. Andersen and I. Chen for discussions and editing the manuscript; A. N. Kettenbach for advice; J. Houston and K. Lowe for FACS; P. Sinojia and E. Rivera-Peraza for preliminary data analyses. Sequencing was performed at the Harwell Center for Biotechnology and images were acquired at the Cell & Tissue Imaging Center, both of which are supported by SJCRH and NCI P30 (CA021765). M.N.D, Y.P., and B.X. are supported by NCI P30 grant (CA21765). J.P. is supported by NIH (RF1 AG064909, RF1AG068581, U54NS110435, and U19AG069701). This work is funded by the American Lebanese Syrian Associated Charities, American Cancer Society (132096-RSG-18-032-01-DDC), and NIH (1R01GM134358-01). The content is solely the responsibility of the authors and does not necessarily represent the official views of the National Institutes of Health. The funders had no role in study design, data collection and analysis, decision to publish, or preparation of the manuscript.

## AUTHOR CONTRIBUTIONS

B.L: most experiments and data analyses. M.N.D., B.X., and Y.F.: bioinformatics analyses. Y.M.: *ATM*-KO generation. S.J.: histone WB and data analyses. Z.F.Y, X.W., K.K., V.P., and J.P.: proteomics data acquisition and analyses. X.Y.: NPC generation. A.S.P.: image quantification. H.P. and F.W.: Supplementary Tables 3 and 4. S.P.M.: design of 53BP1-S25A and S25D mutagenesis. J.C.P.: project design, data analyses, and manuscript writing with inputs from all authors.

## COMPETING INTERESTS

The authors declare no competing interests.

## MATERIALS & METHODS

### Buffers

PBS: 137 mM NaCl, 2.7 mM KCl, 10 mM Na2HPO4, 1.8 mM KH2PO4 (pH 7.4). PBST: PBS with 0.1% Triton X-100. HEPM: 25 mM HEPES (pH 6.9), 10 mM EGTA, 60 mM PIPES, 2 mM MgCl2. Immunofluorescence blocking solution: 1/3 Blocker Casein (ThermoFisher Scientific), 2/3 HEPM with 0.05% TX-100. Buffer A: 10 mM HEPES (pH 7.9), 10 mM KCl, 1.5 mM MgCl2, 0.34 M sucrose, 10% glycerol. Buffer B: 3 mM EDTA, 0.2 mM EGTA. Buffer D: 400 mM KCl, 20 mM HEPES, 0.2 mM EDTA, 20% glycerol. ChIP lysis buffer 3: 10 mM Tris-HCl (pH 8.0), 100 mM NaCl, 1 mM EDTA, 0.5 mM EGTA, 0.1% sodium deoxycholate, 0.5% N-Lauroylsarcosine. ChIP wash buffer: 50mM HEPES (pH 7.5), 500mM LiCl, 1mM EDTA, 1% NP-40, 0.7% Na-deoxycholate. ChIP elution buffer: 50 mM Tris-HCl (pH 8.0), 10mM EDTA, 1% SDS.

### Antibodies

Supplementary Table 5 lists all antibodies and conditions used in this study.

### ESC culture and mutagenesis

H9/WA09 (WiCell) hESCs were grown on Matrigel with reduced growth factors (ThermoFisher Scientific, #35423) in mTeSR1 medium (STEMCELL Technologies, #85850) at 37°C and 5% CO_2_. The 53BP1 knockin cell lines (53BP1 S25A 34-3, 34-4, 79-1, 79-3 and S25D 14-3, 14-15, 14-19, 17) and ATM knockout cell lineS (ATM-KO2, 3, 14, and 43) were generated using CRISPR/Cas9 gene-editing technology. Genome editing reagents were designed and validated in the Center for Advanced Genome Engineering at St. Jude Children’s Research Hospital. Briefly, a chemically modified sgRNA (Synthego) was precomplexed with *SpCas9* protein (St. Jude Protein Production Core) and co-transfected with an ssODN donor template containing the desired modification into H9/WA09 cells via nucleofection (Amaxa P3 primary cell 4D nucleofector X kit L, Lonza) using the manufacturer’s recommended protocol. Transfected cells were sorted (BD FACSAria™ Fusion) onto Matrigel and allowed to grown single-cell clones. Clones were identified via targeted mi-seq using a two-step PCR library setup as previously described (*33*). Samples were demultiplexed using the index sequences, fastq files were generated, and NGS analysis was performed using CRIS.py (*34*). Supplementary Table 6 lists genome editing reagents and associated primers.

### Neural progenitor cell generation and culture

ESCs were seeded onto AggreWell800 plates (STEMCELL Technologies, #34811) and fed with neural induction medium (STEMCELL Technologies, #05835) to form embryoid bodies. On day 5, embryoid bodies were re-plated onto Matrigel-treated 6-well plates in the same media.. On day 17, cells were harvested as NPCs.

### Nuclear extract preparation and Western blotting

ESCs and NPCs were incubated in Buffer A + PI + DTT for 5 min on ice. After centrifugation at 1750 *g* for 2 min at 4 °C, the nuclei pellet was washed in Buffer A and subsequently incubated for ∼25 min in Buffer D + PI + DTT at 4 °C with rotation to obtain the nuclear fraction. Nuclear extracts were separated by SDS–PAGE and transferred onto a nitrocellulose membrane (Bio-Rad). Membranes were blocked with 3% bovine serum albumin (BSA) in HEPM, incubated in primary antibodies (HEPM containing 1% BSA and 0.1% Triton X-100) overnight at 4 °C, washed in PBS-T, incubated in IRDye-conjugated secondary antibodies (LI-COR), and imaged on an Odyssey Fc imaging system (LI-COR). Signals were quantitated with the Image Studio software (version 1.0.14; LI-COR).

### Immunoprecipitation

Antibody was bound to protein A and protein G Dynabeads™ (ThermoFisher 10002D and 10004D) for 2 h at room temperature. Nuclear extract was incubated with the Dynabeads-antibody complex for 5 h at 4 °C, washed with PBST, and eluted with 0.1 M glycine (pH 2.3). Eluates were neutralized with 1/10 volume of 1.5 M Tris buffer (pH 8.8).

### Cortical organoid differentiation

Cortical organoids were generated based on previously published methods with minor modifications (*35, 36*). In brief, hESC lines were expanded and dissociated to single cells using Accutase, seeded onto low-attachment V-bottom 96-well plates (Costar, #7007) at a density of 9,000 cells per well to aggregate into embryoid bodies. The embryoid bodies formation medium (DMEM/F-12 with 20% knockout serum replacement, 3% ESC-quality FBS, 2 mM GlutaMAX, 0.1 mM nonessential amino acids) was supplemented with dorsomorphin (2 μM), WNT inhibitor (IWR1, 3 μM), TGF-β inhibitor (SB431542, 5 μM), and Rho kinase inhibitor (Y-27623, 20 μM). Starting from day 4, embryoid bodies were fed with cortical differentiation medium (Glasgow-MEM, 20% KSR, 0.1 mM NEAA, 1 mM sodium pyruvate, 0.1 mM β-ME, and 1% anti-anti), supplemented with WNT inhibitor (IWR1, 3 μM), TGF-β inhibitor (SB431542, 5 μM), cyclopamine (2.5 μM) and Rho kinase inhibitor (Y-27623, 20 μM). On day 17, embryoid bodies were embedded in Matrigel droplets and transferred onto low-attachment 6-wells and cultured in suspension using DMEM/F-12 supplemented with 1% N2 supplement, 1% lipid concentrate, 2% B27 supplement without vitamin A, and 1% anti-anti under 40% O_2_/5% CO_2_ conditions on shaker. Starting from day 30, medium was changed to 50% DMED/F-12, 50% neurobasal media, 0.5% N2 supplement, 1% GlutaMax, 0.05 mM NEAA, 0.025% human insulin, 0.1 mM β-ME, and 1% anti-anti, supplemented with 2% B27.

### Immunofluorescence

Cells and cryosectioned organoids were blocked with IF blocking solution for 2 h at RT and primary antibodies (diluted in blocking buffer) added and incubated O/N at 4 °C. After 3 washES in PBS-T, fluorescent dye-conjugated secondary antibodies (1:500, Alexa Fluor-CONJUGATED antibodies, ThermoFisher Scientific) were added and incubated for 3 h at RT. Secondary was washed with PBS-T three times and samples were washed and coverslips mounted with Prolong Glass Mounting Reagent (ThermoFisher Scientific) which contains DAPI. Images were acquired with Zeiss LSM780.

### Organoid feature characterization by image analysis

At days 35 and 55, bright-field images of organoids were captured with Axiocam 208 (Zeiss). Area of organoids, area of ventricular zone-like regions, and marker-positive cells were quantified by using the software FIJI: signals-positive cells were identified based on signal and width thresholds. For ventricular zone-like region quantification, inner and outer edges of the regions in the image were manually traced. FIJI was used to quantify area, perimeter, major and minor axes of the inner and outer traces. Mean perimeter and the difference between the major axes of the inner and outer traces were used to estimate the thickness of the structure. Mean Perimeter = (outer perimeter + inner perimeter)/2. MajorAxisDiff = (outer major axis – inner major axis)/2. MinorAxisDiff=(outer minor axis – inner minor axis)/2.

## RNA-seq

Total RNA was extracted with TRIzol reagent (Invitrogen™, #15596026) and Direct-zol™ RNA Microprep (Zymo Research, # R2062) by following manufacturer’s instructions. DNA digestion with DNase I was performed during RNA extraction. Paired-end 100-cycle sequencing was performed on NovaSeq6000 sequencer by following the manufacturer’s instructions (Illumina). Raw reads were first trimmed using TrimGalore (version 0.6.3) available at: https://www.bioinformatics.babraham.ac.uk/projects/trim_galore/, with parameters ‘--paired –– retain_unpaired’. Filtered reads were then mapped to the *Homo sapiens* reference genome (GRCh38 + Gencode-v31) using STAR (version 2.7.9a) (*37*). Gene-level read quantification was done using RSEM (version 1.3.1) (*38*). To identify the differentially expressed genes between control and experimental samples, the variation in the library size between samples was first normalized by trimmed mean of *M* values (TMM) and genes with CPM < 1 in all samples were eliminated. Then, the normalized data were applied to linear modeling with the voom from the limma R package (*39*). GSEA was performed against using the MSigDB database (version 7.1), and differentially expressed genes were ranked based on log_2_(FC) (*40, 41*).

## Protein extraction, digestion, and Tandem-Mass-Tag (TMT) labeling

Organoids were harvested on day 35 and the Matrigel droplets were eliminated by multiple ice-cold PBS washes. The organoid pellet was extracted in the lysis buffer (50 mM HEPES, pH 8.5, 8 M urea, and 0.5% sodium deoxycholate, 100 μl buffer per 10 mg tissue) with 1× PhosSTOP phosphatase inhibitor cocktail (Sigma-Aldrich). Protein concentration was estimated by a Coomassie stained short gel with bovine serum albumin (BSA) as a standard. About 600µg each of protein samples was digested with LysC (Wako) at an enzyme-to-substrate ratio of 1:100 (w/w) for 2h at room temperature (RT) in the presence of 1mM DTT. The samples were then diluted to a final 2M Urea concentration with 50mM HEPES (pH 8.5), and digested with Trypsin (Promega) at an enzyme-to-substrate ratio of 1:50 (w/w) for 3h. The peptides were reduced by adding 1mM DTT for 30 min at RT followed by alkylation with 10mM iodoacetamide for 30 min in the dark at RT. The unreacted iodoacetamide was quenched with 30mM DTT for 30 min. Finally, the digestion was terminated and acidified by adding trifluoroacetic acid to 1%, peptides desalted using Sep-Pak C18 cartridge (Waters), and dried by speed vac. The purified peptides were resuspended in 50mM HEPES (pH 8.5), labeled with 16-plex Tandem Mass Tag (TMTpro) reagents (Thermo Scientific) following the manufacturer’s recommendation. The TMT labeled samples were mixed equally, desalted using Sep-Pak C18 cartridge (Waters), and dried by speed vac.

## Offline fractionation and two-dimensional liquid chromatography-tandem mass spectrometry (LC/LC-MS/MS)

The dried TMT mix was resuspended and fractionated on an offline HPLC (Agilent 1220) using basic pH reverse phase liquid chromatography (pH 8.0, XBridge C18 column, 4.6 mm x 25 cm, 3.5 μm particle size, Waters). A total of 160 one-minute fractions were collected and concatenated to 80 fractions. 10% of these 80 fractions was used for whole proteome analysis. The remaining 90% of the 80 fractions were concatenated to 20 fractions for phophopeptide enrichment. Phosphopeptide enrichment was performed according to a previously published protocol (*42*). The phosphopeptide enrichment eluents and the total proteome fractions were dried and resuspended in 5% formic acid and analyzed by acidic pH reverse phase LC-MS/MS analysis. The peptide samples were loaded on a nanoscale capillary reverse phase C18 column (New objective, 75 μm ID x ∼15 cm, 1.9 μm C18 resin from Dr. Maisch GmbH) by a HPLC system (Thermo Ultimate 3000) and eluted by either a 125 min gradient (phospho fractions) or 110 min gradient for total proteome fractions. The eluted peptides were ionized by electrospray ionization, and detected by an inline Orbitrap Fusion mass spectrometer (Thermo Scientific). For total proteome fractions, the mass spectrometer is operated in data-dependent mode with a survey scan in Orbitrap (60,000 resolution, 2 × 10^5^ AGC target and 50 ms maximal ion time) and MS/MS high resolution scans (60,000 resolution, 1 × 10^5^ AGC target, 150 ms maximal ion time, 36.5 HCD normalized collision energy, 1 *m/z* isolation window, and 15 s dynamic exclusion). For phosphoproteome fractions, the mass spectrometer is operated in data-dependent mode with a survey scan in Orbitrap (60,000 resolution, 3 × 10^5^ AGC target and 50 ms maximal ion time) and MS/MS high resolution scans (60,000 resolution, 1 × 10^5^ AGC target, 150 ms maximal ion time, 36.5 HCD normalized collision energy, 1 *m/z* isolation window, and 10 s dynamic exclusion).

## Identification of proteins and phosphopeptides

The MS/MS raw data were processed by a tag-based hybrid search engine JUMP (*43*). The data was searched against the UniProt human database (168,305 protein entries; downloaded in April 2020) concatenated with a reversed decoy database for evaluating FDR. Searches were performed using a 15 ppm mass tolerance for fragment ions, fully tryptic restriction with two maximal missed cleavages, three maximal modification sites, and the assignment of *b*, and *y* ions. TMT tags on Lysine residues and N-termini (+304.2071453 Da) were used for static modifications and Met oxidation (+15.99492 Da) was considered as a dynamic modification. Phosphorylation (+79.96633 Da) was considered as a dynamic modification for STY residues. Putative peptide spectral matches (PSMs) were filtered by mass accuracy and then grouped by precursor ion charge state and filtered by JUMP-based matching scores (Jscore and ΔJn) to reduce FDR below 1% for proteins during the whole proteome analysis or 1% for phosphopeptides during the phosphoproteome analysis. Phosphosites were further evaluated by JUMPl program using the concept of the phosphoRS algorithm (*44*) to calculate phosphosite localization scores (Lscore, 0%–100%) for each PSM.

## Quantification of proteins and phosphopeptides

TMT reporter ion intensities of each PSM were extracted and corrected based on isotopic distribution of each labeling reagent. Those PSMs with very low intensities (e.g., minimum intensity of 1,000 and median intensity of 5,000) were excluded for quantification. Sample loading bias was mitigated by normalization with the trimmed median intensity of all PSMs. Protein or phosphopeptide relative intensities were calculated by dividing the intensity of each channel by the mean intensity. Protein or phosphopeptide absolute intensities were computed by multiplying the relative intensities by the grand-mean of three most highly abundant PSMs.

## Differential expression analysis of proteins and phosphopeptides

Differentially expressed proteins between the two strains and two different doses were identified by the limma R package (*45*). The Benjamini-Hochberg method was used to control multiple-testing correction, and proteins with an adjusted *p* value of < 0.05 and log2 fold change of > 1.5 were defined as differentially expressed.

## Pathway enrichment analysis for proteomics data

Pathway enrichment analysis was carried out to infer functional groups of proteins that were enriched in a given dataset. The four common pathway databases were used, including Gene Ontology (GO), KEGG, Hallmark, and Reactome. The analysis was performed using Fisher’s exact test with the BH correction for multiple testing. A cutoff of adjusted *p* value < 0.2 was used to identify significantly enriched pathways.

## Estimation of kinase activity

Kinase activity was inferred based on known substrates in the PhosphoSitePlus database (*46*) using the IKAP algorithm (*24*). The phosphoproteome data were normalized against the whole proteome. We performed 100 times of calculations to overcome the potential problem of local optimization.

## Chromatin immunoprecipitation

Cells were harvested in PBS. Cytoplasmic fractions were extracted using buffer A with 1× protease inhibitors and 1 mM DTT. Nuclear pellets were cross-linked by 1.1% formaldehyde in buffer B with 1× protease inhibitors and 1 mM DTT; washed; and lysed in lysis buffer 3 with 1× protease inhibitors, 1 mM DTT, and 1 mM PMSF. The fixed and lysed nuclear extract was sonicated with Bioruptor^®^ Pico (Diagenode) 10 times for 15 s each, with 45-s intervals. Chromatin was added to Dynabeads™ (Life Technologies) prebound with 4 µg of antibodies for overnight incubation. After incubation, beads were washed and immunoprecipitates were eluted. DNA from eluates was recovered by the GeneJET FFPE DNA purification kit (ThermoFisher Scientific, #K0882). DNA libraries were generated using the NEBNext™ Ultra DNA Library Prep kit (NEB, #E7370S) and sequenced at the St. Jude Hartwell Center.

## Analysis of chromatin immunoprecipitation-sequencing

50bp single-end reads were obtained and aligned to human genome hg19 by BWA (version 0.7.170.7.12, default parameter). Duplicated reads were marked by the bamsormadup from the biobambam tool (version 2.0.87) available at https://www.sanger.ac.uk/tool/biobambam/. Uniquely mapped reads were kept by samtools (parameter “-q 1 –F 1804,” version 1.14). Fragments < 2000 bp were kept for peak calling and bigwig files were generated for visualization. SICER (*27*) and macs2 (*28*) were both used for peak calling, to identify both the narrow and broad peak correctly. With SICER, we assigned peaks that were at the top 1 percentile as the high-confidence peaks and the top 5 percentile as the low-confidence peaks. Two sets of peaks were generated: Strong peaks called with parameter ‘FDR < 0.05’ by at least one method (macs2 or SICER) and weak peaks called with parameter ‘FDR < 0.5’ by at least one method (macs2 or SICER). Peaks were considered reproducible if they were supported by a strong peak from all replicates or at least one strong peak and a weak peak in the other replicates. For downstream analyses, heatmaps were generated by deepTools (*47*) and gene ontology was performed with Enrichr (*48, 49*) and GSEA, in addition to custom R scripts. For differential peak analysis, peaks from two replicates were merged and counted for number of overlapping extended reads for each sample (bedtools v2.24.0) (*50*). Then we detected the differential peaks by the empirical Bayes method (eBayes function from the limma R package) (*39*). For downstream analyses, heatmaps were generated by deepTools (v3.5.0) (*51*). Peaks were annotated based on Gencode following this priority: “Promoter.Up”: if they fall within TSS – 2kb, “Promoter.Down”: if they fall within TSS – 2kb, “Exonic” or “intronic”: if they fall within an exon or intron of any isoform, “TES peaks”: if they fall within TES ± 2kb, “distal5” or “distal3” if they are with 50kb upstream of TSS or 50kb downstream of TES, respectively, and they are classified as “intergenic” if they do not fit in any of the previous categories.

---

**Supplementary Figure 1.**
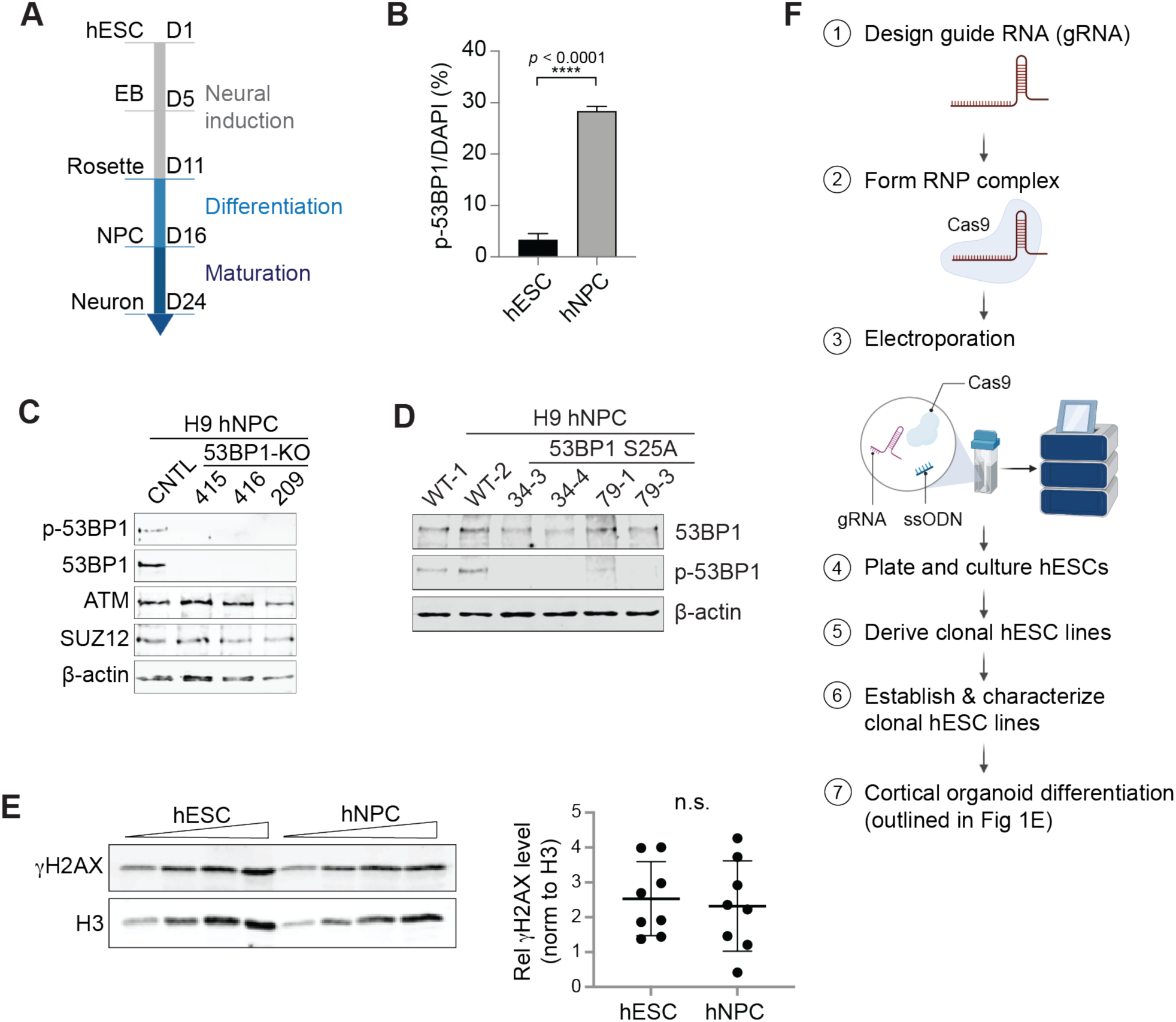
Characterization of 53BP1-pS25 and genome editing of hESCs. (A) Schematic diagram of neural differentiation of hESCs: neural induction, differentiation, and maturation media to form EBs (embryoid bodies), rosettes, NPCs, and neurons. (B) Quantification of 53BP1-pS25-positive hESCs or hNPCs. Data are presented as the mean ± SEM, with *p*<0.0001. (C) WB analysis of control cells and 53BP1-KO clones 415, 416, and 209, which are clones KO1, KO2, and KO3 in Yang, Xu et al. 2019. (D) WB analysis of control and 53BP1-S25A hNPCs. The S25A mutation prohibits phosphorylation. (E) WB analysis of hESCs and hNPCs and quantification. (F) Schematic diagram of genome editing in hESCs. Guide RNA 6 were complexed with Cas9 proteins and used along single-stranded nucleotide donors to transfect hESCs. Individual clones from transfection were cultured, sequenced by mi-seq across the targeted *53BP1* locus, and established as >99% pure clonal lines.

**Supplementary Figure 2.**
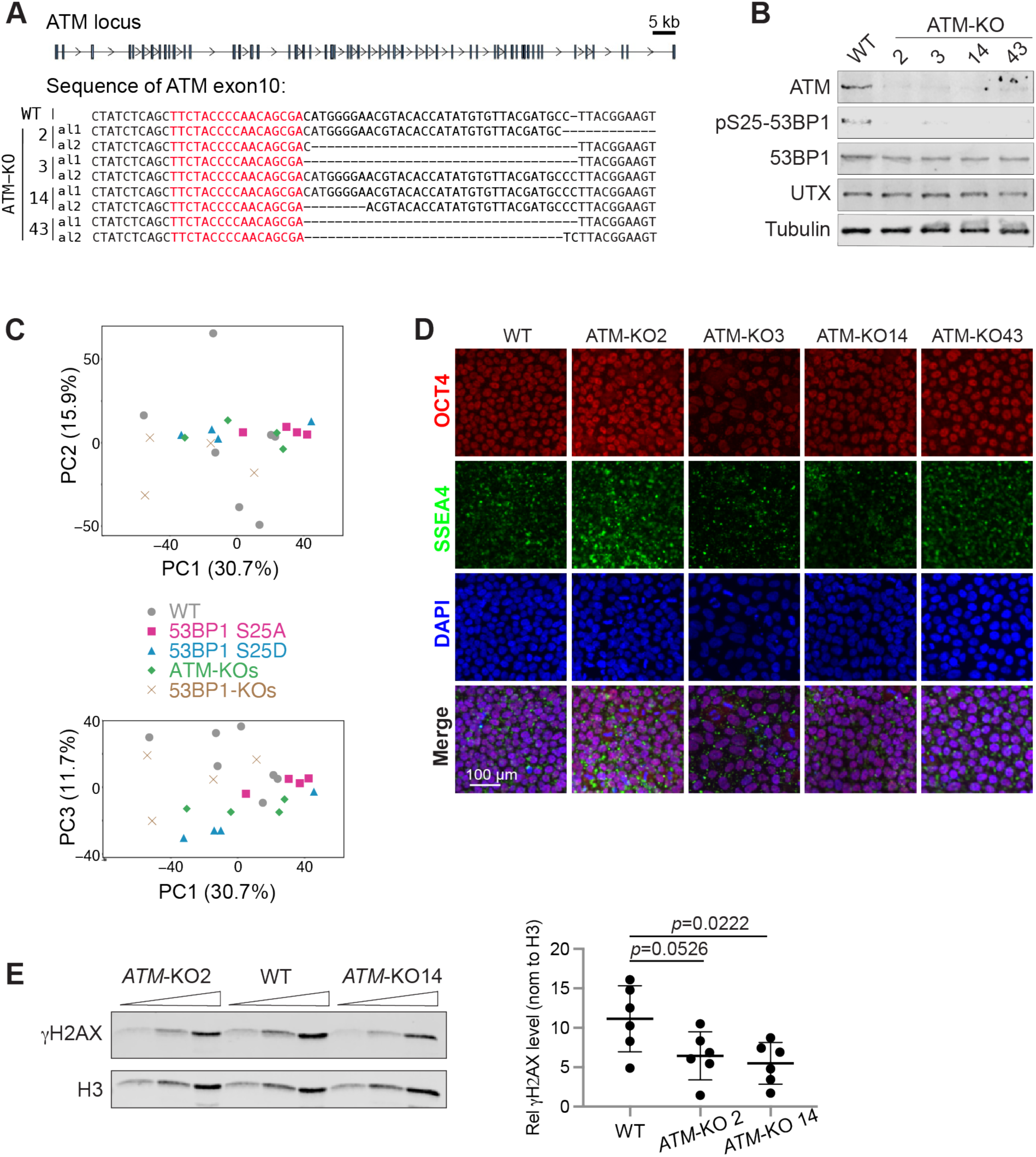
Generation and analyses of ATM-KO hESCs and cortical organoids. (A) Alignment of WT and *ATM*-KO mutation sequences on 2 alleles (al) in the *ATM* locus. Red indicates the gRNA sequence. (B) WB analysis of WT and 4 *ATM*-KO hNPCs. (C) Principal component analysis showed the intermixing and similar RNA-seq profiles from hESCs of 7 WT, 4 53BP1-S25A, 4 53BP1-S25D, 4 ATM-KO, and 4 53BP1-KO lines. (D) Immunofluorescence showed similar expression of OCT4 and SSEA4 proteins in control and *ATM*-KO hESCs. Bar, 100 μm. (E) WB analysis of WT and 2 *ATM*-KO hNPCs. Quantification suggests reduction of γH2AX in *ATM*-KO hNPCs. Welch’s t test was used to perform pairwise comparisons of WT and *ATM*-KO.

**Supplementary Figure 3.**
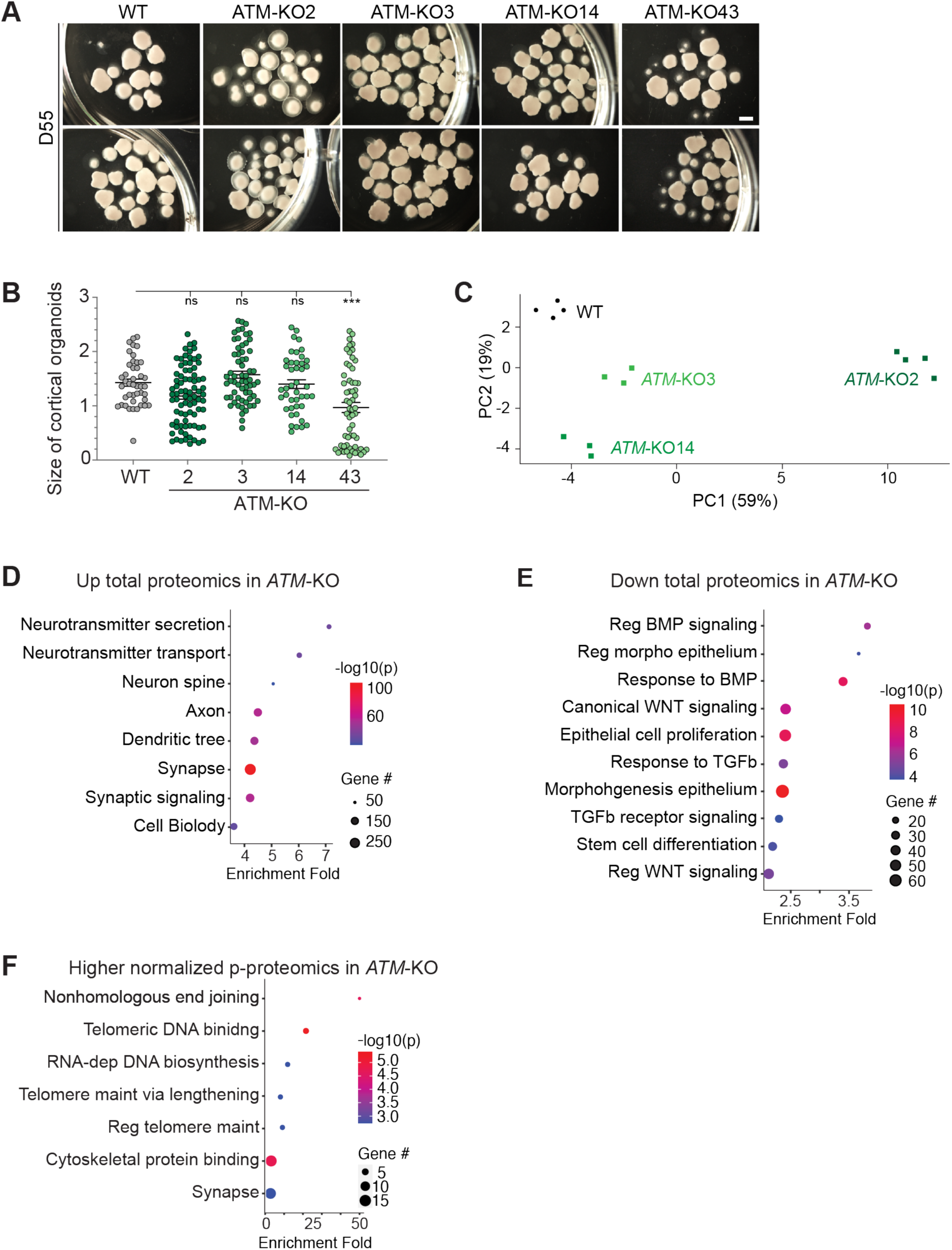
Characterization of D35 *ATM*-KO cortical organoids. (A) Bright-field images of cortical organoids formed by *ATM*-KO2, 3, 14, 43, and WT control at day 55 of differentiation. Bar, 1.5 mm. The (B) size of cortical organoids was compared between groups by one-way ANOVA with Dunnett’s multiple comparisons test to yield n.s. and *** indicating not significant and *p*<0.001, respectively. n = 13 organoids/group. (C) Principal component analysis of proteomics data of D35 WT and *ATM*-KO cortical organoids. GSEA terms that are highly enriched in significantly (C) higher and (D) lower total proteins in D35 *ATM*-KO versus WT cortical organoids. (F) GSEA terms that are highly enriched in significantly higher phospho-proteins, which were normalized to total proteomics, in D35 *ATM*-KO versus WT cortical organoids.

**Supplementary Figure 4.**
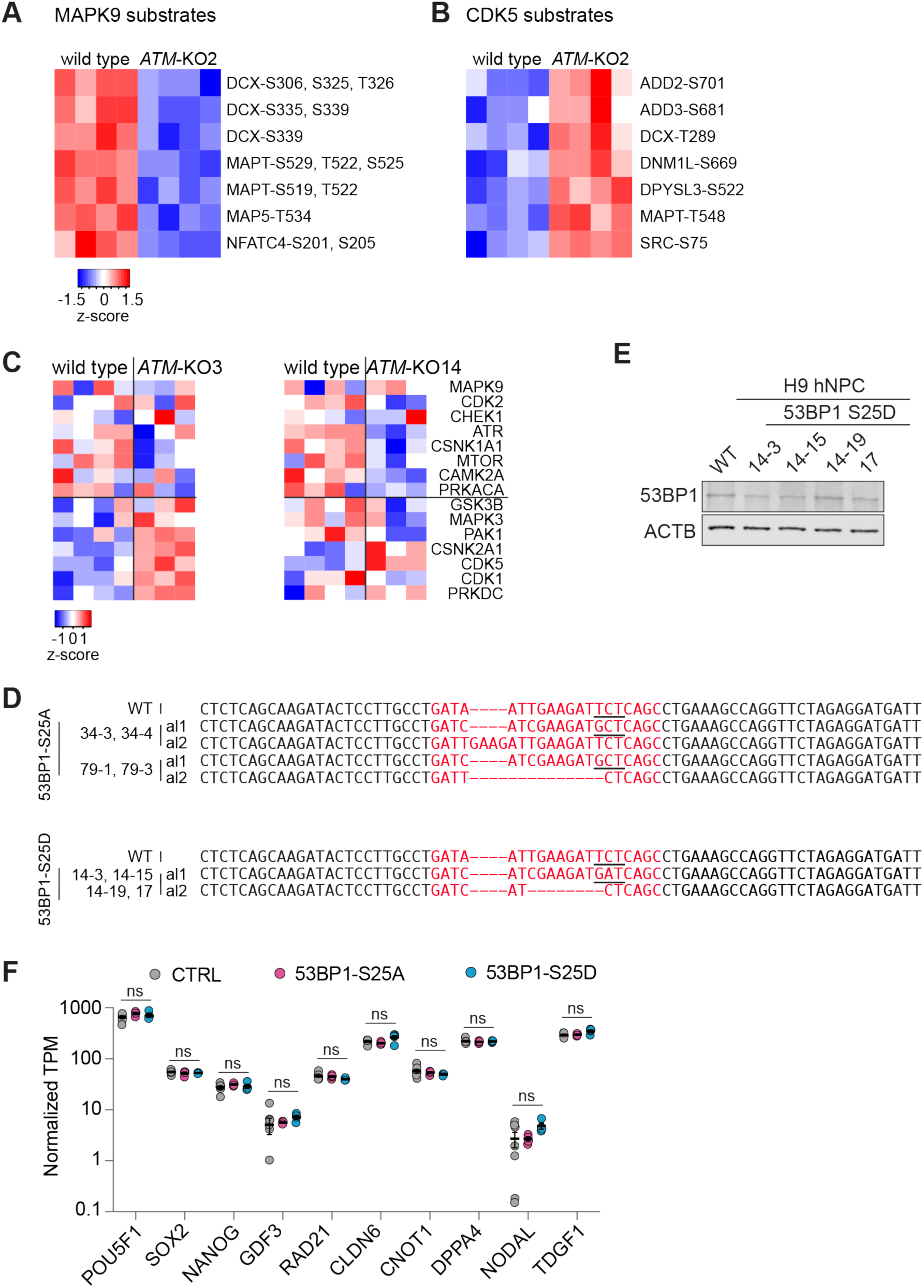
Kinase activities in cortical organoids and characterization of the 53BP1-S25A and –S25D hESCs. Heatmaps showing relative phosphorylation levels of (A) 7 MAPK9 substrates that are significantly lower and (B) 7 CDK5 substrates that are significantly higher in D35 *ATM*-KO versus WT cortical organoids. (C) Heatmaps showing activity of selected protein kinases between ATM-KO3, ATM-KO4, and wild type cell lines. (D) Alignment of WT and 53BP1-S25A and S25D mutation sequences on 2 alleles (al). Red indicates the gRNA sequence. Underline indicates codon encoding the wild-type serine 25, mutant alanine, or mutant aspartic acid. (E) WB analysis of control and 53BP1-S25D hNPCs, which have comparable levels of 53BP1 protein. (F) Transcripts per million values of 10 pluripotency genes were used for comparison to show that control, 53BP1-S25A, and 53BP1-S25D hESCs did not differ in pluripotency.

**Supplementary Figure 5.**
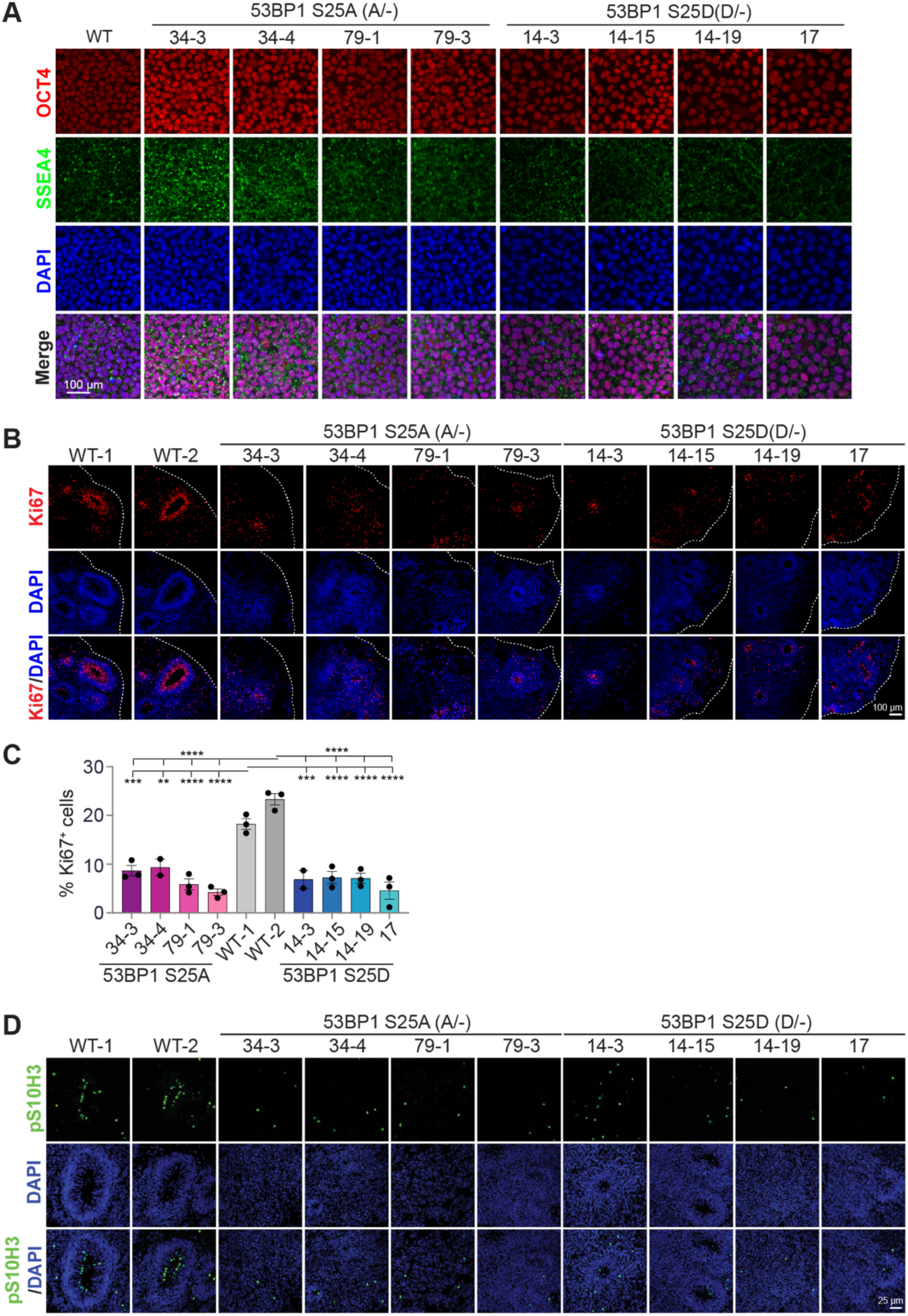
Characterization of the 53BP1-S25A and –S25D hESCs and cortical organoids. (A) Immunofluorescence showed similar expression of OCT4 and SSEA4 proteins in WT, 53BP1-S25A, and 53BP1-S25D hESCs. Bar, 100 μm. Immunofluorescence of (B) KI67 and (D) H3-pS10 in cryosections of cortical organoids at day 35 of differentiation. Bar, 100 μm. (C) Quantification of KI67-positive cells in D35 cortical organoids. Data points represent single organoids. The mean ± SEM values were compared by one-way ANOVA with Dunnett’s multiple comparisons test to yield ****, ***, and ** indicating p< 0.0001, 0.001, and 0.01, respectively. n = 3 organoids/group.

**Supplementary Figure 6.**
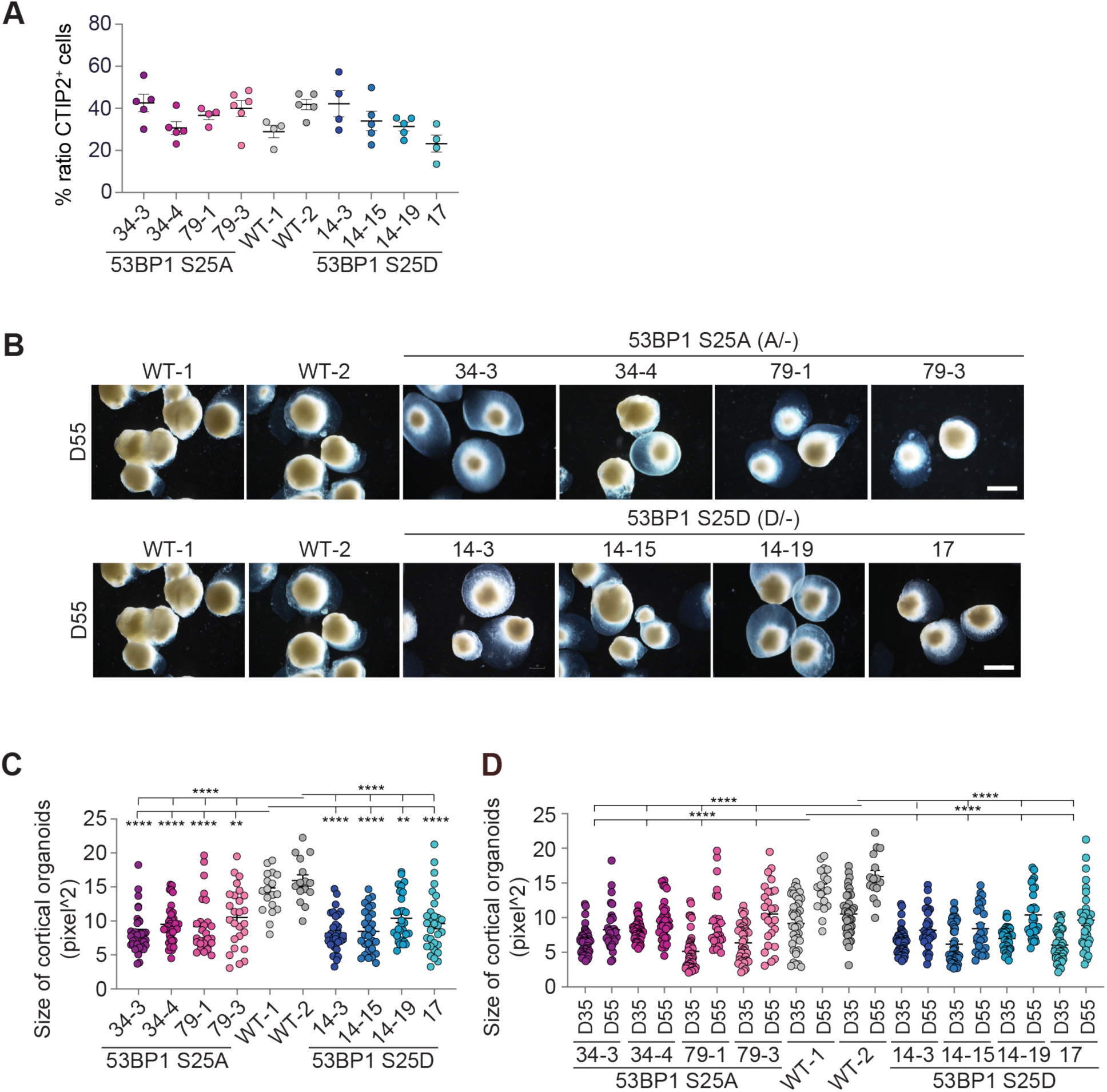
Characterization of 53BP1-S25A and –S25D cortical organoids. (A) Quantification of CTIP2-positive cells in cortical organoids at day 35 of differentiation. Data points represent single organoids. The mean ± SEM values were compared by one-way ANOVA with Dunnett’s multiple comparisons test. (B) Bright-field images of cortical organoids formed by cell lines 53BP1-S25A 34-3, 34-4, 79-1, 79-3 and S25D 14-3, 14-15, 14-19, 17, and 2 WT control at day 55 of differentiation. Bar, 1.5 mm. Blue transparent structures around organoids are Matrigel embedment. At day 55 of differentiation, the (C) size and (D) growth (comparing organoids at days 35 and 55) of cortical organoids were compared between groups. Data points represent single organoids. The mean ± SEM values were compared by one-way ANOVA with Dunnett’s multiple comparisons test to yield **** and ** indicating p< 0.0001 and 0.01, respectively. n = 15 – 36 organoids/group.

**Supplementary Figure 7.**
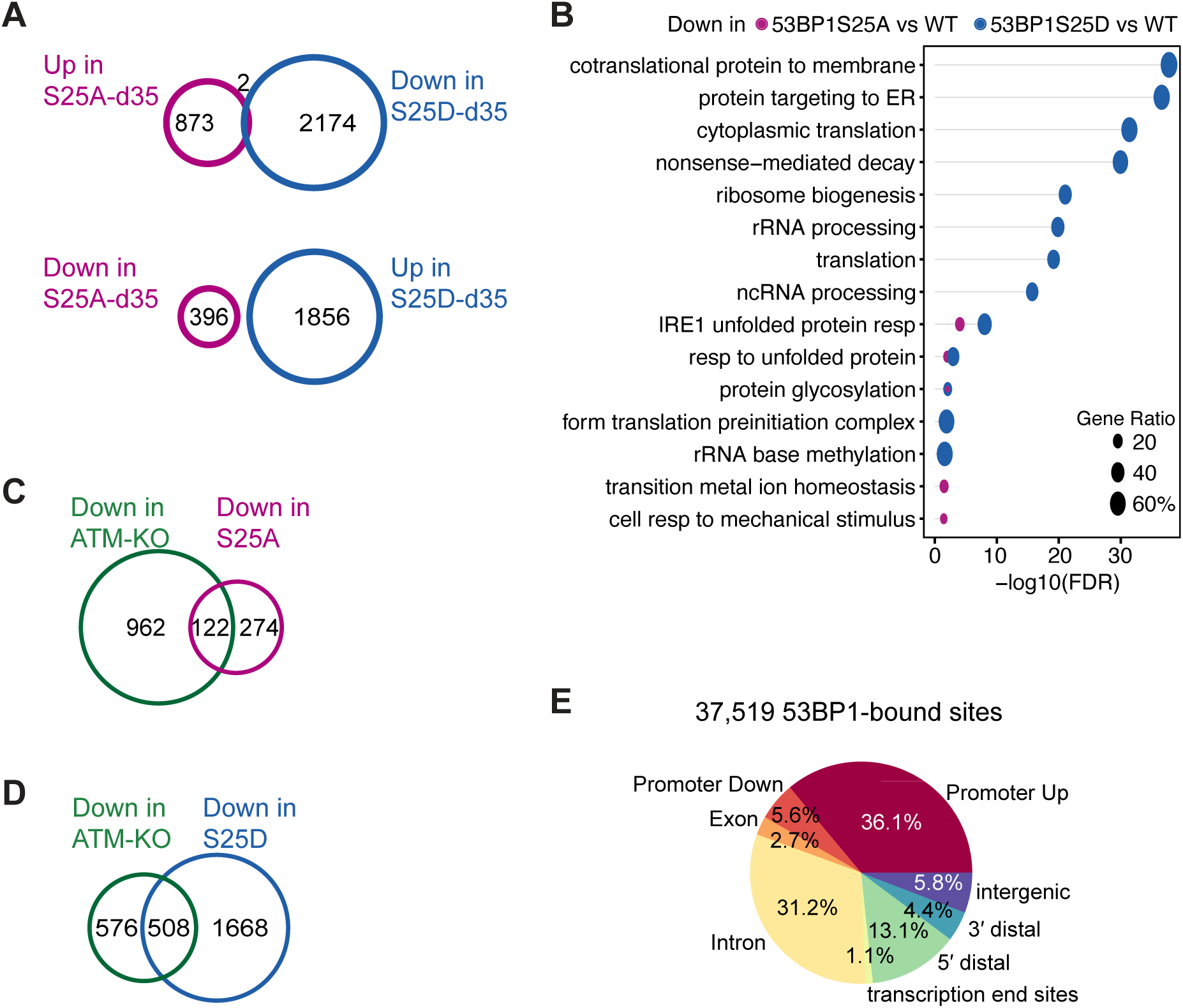
Molecular characterization of cortical organoids and 53BP1 ChIP-seq. (A) Two genes overlapped between upregulated genes in 53BP1-S25A versus WT and downregulated genes in 53BP1-S25D versus WT cortical organoids. No gene overlapped between downregulated genes in 53BP1-S25A versus WT and upregulated genes in 53BP1-S25D versus WT cortical organoids. (B) Downregulated GSEA terms between 53BP1-S25A versus WT and 53BP1-S25D versus WT were not highly overlapped. Ten GSEA terms were specific to 53BP1-S25D versus WT. Venn diagrams depict overlaps between downregulated genes in *ATM*-KO with 53BP1-(C) S25A or (D) S25D cortical organoids. (E) Proportions of 53BP1 binding to genomic features.

**Supplementary Table 1.** List of phospho-proteins, normalized to total protein levels, that were significantly lower in D35 *ATM*-KO versus WT cortical organoids.

**Supplementary Table 2.** Normalized (to total proteome) levels of phosphor-peptide substrates of MAPK9 and CDK5 in D35 WT and *ATM*-KO cortical organoids.

**Supplementary Table S3.**
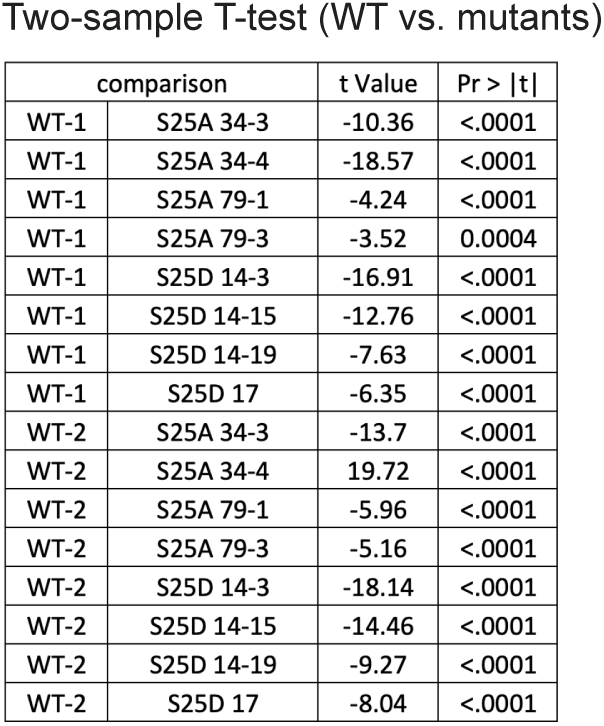
Two-sample *t*-test examines the sizes of cortical organoids that change between day 35 and day 55 of differentiation. Data from wild-type and 53BP1 mutants are compared pair-wise by using the two-sample t-test. The sizes of organoids are significantly different between each comparison pair (all p < 0.05).

| comparison |  | t Value | Pr > \|t\| |
| --- | --- | --- | --- |
| WT-1 | S25A 34-3 | -10.36 | <.0001 |
| WT-1 | S25A 34-4 | -18.57 | <.0001 |
| WT-1 | S25A 79-1 | -4.24 | <.0001 |
| WT-1 | S25A 79-3 | -3.52 | 0.0004 |
| WT-1 | S25D 14-3 | -16.91 | <.0001 |
| WT-1 | S25D 14-15 | -12.76 | <.0001 |
| WT-1 | S25D 14-19 | -7.63 | <.0001 |
| WT-1 | S25D 17 | -6.35 | <.0001 |
| WT-2 | S25A 34-3 | -13.7 | <.0001 |
| WT-2 | S25A 34-4 | 19.72 | <.0001 |
| WT-2 | S25A 79-1 | -5.96 | <.0001 |
| WT-2 | S25A 79-3 | -5.16 | <.0001 |
| WT-2 | S25D 14-3 | -18.14 | <.0001 |
| WT-2 | S25D 14-15 | -14.46 | <.0001 |
| WT-2 | S25D 14-19 | -9.27 | <.0001 |
| WT-2 | S25D 17 | -8.04 | <.0001 |

**Supplementary Table S4.** The changes in organoid size at days 35 and 55 of differentiation were compared to yield Supplementary Figure 3C. This table lists the calculation for different combinations of data and the descriptive statistics.

| Type | n pairs<br>(D55-D35) | n<br>organoids@<br>D35 | n<br>organoids@<br>D55 | Mean | STDDEV | Min | Max |
| --- | --- | --- | --- | --- | --- | --- | --- |
| WT-1 | 720 | 40 | 18 | 4978.41 | 4652.28 | -7099 | 16018 |
| WT-2 | 585 | 39 | 15 | 5407.41 | 4475.87 | -7464 | 19093 |
| S25A 34-3 | 1692 | 47 | 36 | 2993.29 | 3351.85 | -5833 | 16148 |
| S25A 34-4 | 1505 | 43 | 35 | 1427.69 | 3125.62 | -7267 | 9852 |
| S25A 79-1 | 1204 | 43 | 28 | 4029.41 | 4813.68 | -7445 | 17566 |
| S25A 79-3 | 1066 | 41 | 26 | 4157.93 | 5089.87 | -8628 | 17365 |
| S25D 14-3 | 1408 | 44 | 32 | 1693.88 | 3281.67 | -8699 | 10981 |
| S25D 14-15 | 1232 | 44 | 28 | 2286.89 | 4212.01 | -8321 | 12011 |
| S25D 14-19 | 1247 | 43 | 29 | 3422.64 | 3795.38 | -4957 | 13397 |
| S25D 17 | 1333 | 43 | 31 | 3577.88 | 4830.71 | -7145 | 19121 |

**Supplementary Table 5.**

| Primary Antibodies | Company | Cat. No. | Experimental Condition |
| --- | --- | --- | --- |
| 53BP1-pS25 | Abcam | ab70323 | WB (1:1000) |
| 53BP1 | Novus Biological | NB100-304 | WB (1:1000) |
| UTX | Millipore /<br>Cell Signaling Technology | ABE409 /<br>33510 | WB (1:1000)<br>WB (1:1000) |
| PAX6 | BioLegend | 901301 | IF (1:200) |
| CTIP2 | Abcam | Ab18465 | IF (1:500) |
| KI67 | Cell Signaling Technology | 9129S | IF (1:100) |
| β-actin | Sigma Aldrich | A1978 | WB (1:1000) |
| ATM | Abcam | Ab17995 | WB (1:1000) |
| SUZ12 | Cell signaling Technology | 3737S | WB (1:1000) |
| OCT4 | Cell signaling Technology | 2840S | IF (1:200) |
| SSEA4 | STEMCELL Technologies | 60062AD | IF (1:100) |
| pS10H3 | Cell Signaling Technology | 3465S | IF (1:1000) |

**Supplementary Table 6.**
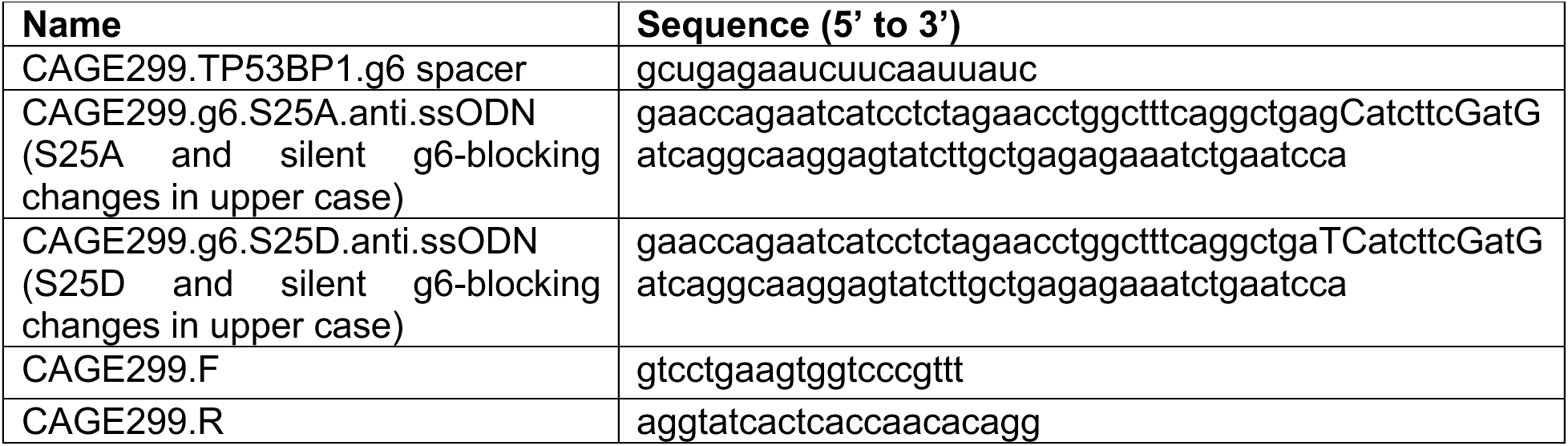

## Notes

### Competing Interest Statement

The authors have declared no competing interest.

